# Dendritic inhibition by Shh signaling-dependent stellate cell pool is critical for motor learning

**DOI:** 10.1101/2021.04.15.439999

**Authors:** Wen Li, Lei Chen, Jonathan T. Fleming, Emily Brignola, Kirill Zavalin, Andre H. Lagrange, Tonia S. Rex, Shane A. Heiney, Gregory J. Wojaczynski, Javier F. Medina, Chin Chiang

**Author notes:** Address all correspondence to: Chin Chiang. These authors contributed equally (Wen Li, Lei Chen).

## Abstract

Cerebellar inhibitory interneurons are important regulators of neural circuit activity for diverse motor and non-motor functions. The molecular layer interneurons (MLI), consisting of basket cells (BCs) and stellate cells (SCs), provide dendritic and somatic inhibitory synapses onto Purkinje cells, respectively. They are sequentially generated in an inside-out pattern from Pax2+ immature interneurons which migrate from the prospective white matter to the ML of the cortex. However, little is known as to how MLI subtype identities and pool sizes are determined, nor are their contributions to motor learning well understood. Here, we show that GABAergic progenitors fated to generate both BCs and SCs respond to the Shh signal. Conditional abrogation of Shh signaling inhibited proliferation of GABAergic progenitors and reduced the number of Pax2^+^ cells, whereas persistent Shh pathway activation increased their numbers. These changes, however, did not affect early-born BC numbers but selectively altered the SC pool size. Moreover, genetic depletion of GABAergic progenitors when BCs are actively generated also resulted in a specific reduction of SCs, suggesting that the specification of MLI subtypes is independent of Shh signaling and their birth order and likely occurs after Pax2^+^ cells settle into their laminar positions in an inside-out sequence. Mutant mice with reduced SC numbers displayed decreased dendritic inhibitory synapses and neurotransmission onto Purkinje cells, resulting in an impaired acquisition of eyeblink conditioning. These findings also reveal an essential role of Shh signaling-dependent SCs in regulating inhibitory dendritic synapses and motor learning.

## INTRODUCTION

The cerebellum plays an essential role in fine motor learning characterized by an adaptive process that involves recurring error-evoked learning to maintain optimum motor performance (Raymond & Medina, 2018). The circuit that enables this mode of learning is comprised of different classes of inhibitory and excitatory neurons that are generated during embryonic and postnatal development from spatially distinct progenitors (Palay & Chan-Palay, 1974a; Zhang & Goldman, 1996; Wang *et al*, 2005; Machold & Fishell, 2005; Hoshino *et al*, 2005). However, it remains unclear how their identities and numbers are specified to generate a functional circuit.

Purkinje cells (PCs) are the sole projection neurons in the cerebellar cortex, with a dendritic plane in the molecular layer where they integrate excitatory and inhibitory input from different sources (Beckinghausen & Sillitoe, 2018). The principal excitatory neurons are granule cells with their axons extending into the molecular layer as parallel fibers that provide excitatory input to PCs. Additionally, PCs receive excitatory input from two classes of extracerebellar afferent projections; climbing fibers of the inferior olive nuclei terminate in the molecular layer and contact PCs, and mossy fibers (from the brainstem and elsewhere) synapse with granule cells and thus influence many PCs at once. The balance to the excitatory neurotransmission is provided largely by GABAergic inhibitory inputs from molecular layer interneurons (MLIs) that consist of basket cells (BCs) and stellate cells (SCs) (Sotelo, 2015). BCs are located in the inner one-third of the ML, and their axons form perineuronal nests or “baskets” as well as specialized structures known as pinceau around the PC soma/axon initial segment (Somogyu & Hámori, 1976). In contrast, SCs occupy the rest of the ML and make direct contact with distal PC dendrites. Despite the apparent morphological and positional differences between BCs and SCs, the precise location and the boundary between these two cell types have not been well defined until the recent identification of Ret, a receptor tyrosine kinase, whose expression is restricted to BCs (Sergaki *et al*, 2017).

Cerebellar inhibitory neurons are generated from spatially distinct germinal zones. In mice, the early-born neurons, including PCs, Golgi (granule layer interneuron), and Lugaro cells, are generated from the ventricular zone (VZ) of the fourth ventricle during early to mid-embryonic stages (Hoshino *et al*, 2005; Sudarov *et al*, 2011). The late-born MLIs are generated sequentially but in an overlapping manner from the prospective white matter (PWM) during the late embryonic to postnatal development, with BCs emerging first and followed by SCs (Zhang & Goldman, 1996). Recent studies have identified stem cell-like astroglia in the PWM as the source of transient amplifying GABAergic progenitors marked by Ptf1a (Fleming *et al*, 2013), a bHLH transcription factor whose function is required for the specification of all GABAergic lineages, including Pax2^+^ immature interneuron (Hoshino *et al*, 2005; Pascual *et al*, 2007). Pax2^+^ cells migrate from PWM to ML before terminally differentiating into BCs and SCs (Weisheit *et al*, 2006). However, it remains unclear as to how MLI subtype identities are determined.

The Shh signaling pathway has been shown to play critical roles during cerebellar development. It is required for the rapid expansion of granule cells by promoting the proliferation of granule cell precursors (Dahmane & Altaba, 1999; Wallace, 1999; Wechsler-Reya & Scott, 1999). Additionally, Shh signaling is transiently required to maintain the proliferative capacity of multipotent radial glial cells in the VZ and astroglia in the PWM (Huang *et al*, 2009; Fleming *et al*, 2013). Unlike VZ, Ptf1a+ progenitors are proliferative and exhibit Shh pathway activity in the PWM (Fleming *et al*, 2013). However, the significance of this activity in the regulation of MLI pool size and cerebellar function has not been determined. While recent studies have shown that MLIs play an important role in cerebellar-dependent motor learning (Brinke *et al*, 2015; Sergaki *et al*, 2017), it remains to be determined as to what extent each MLI subtype contributes to motor learning. In this study, we investigated the role of Shh signaling and its contribution to MLI subtype allocations and cerebellar-dependent motor learning.

## RESULTS

### Shh signaling is transiently activated in Ptf1a+ progenitors in the PWM

To better define Shh signaling in GABAergic progenitors, we used β-galactosidase (β-gal) expression from *Gli*^*nlacZ*^ mice (Bai *et al*, 2002) to assess Shh pathway responsiveness among Ptf1a^+^ cells at different postnatal stages. We found that Shh signaling is transiently activated in a subset of Ptf1a+ cells from P1 to P8 (Figure 1A), while contemporaneous populations of Ptf1a^+^ cells in the brain/ brainstem were negative for the signal. At P1, the β-Gal+ and Ptf1a+ cells account for ∼19% of Ptf1a+ progenitors in the PWM, and this number increased to ∼26% at P3 (Figure 1B and 1C). However, at P6, the percentage of double-positive cells declined to ∼14%, but no statistically significant changes were observed for Ptf1a+ progenitors (Figure 1B and 1C). By P8, a very small population of Ptf1a+ progenitors were positive for β-gal expression (Figure 1C). Thus, the Shh pathway is transiently activated in a subset of Ptf1a^+^ progenitors.

**Figure 1.**
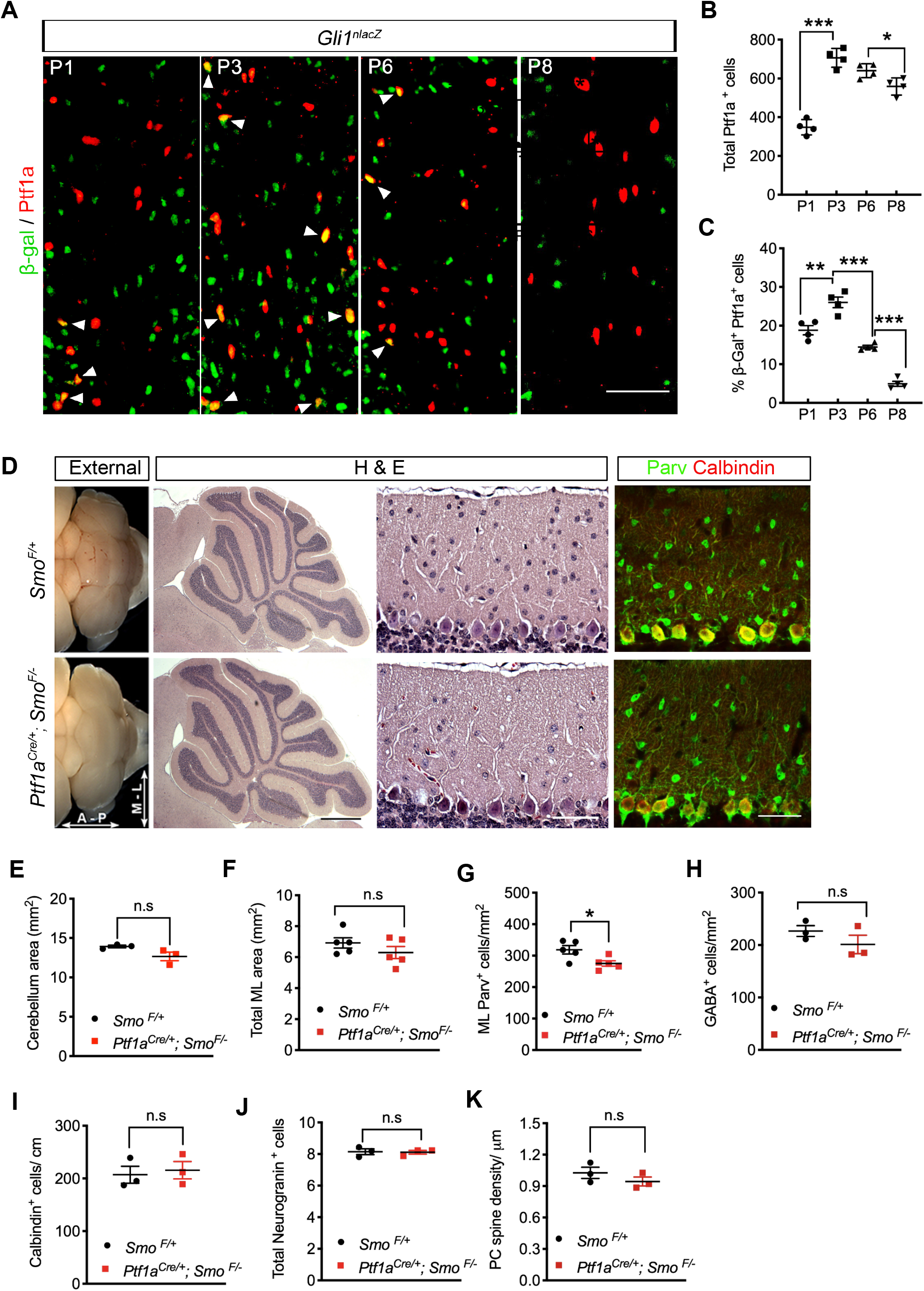
Shh signaling is transiently activated in Ptf1a+ progenitors and is required for MLI expansion. (A) Midsagittal sections of *Gli1*^*nlacZ*^ mice showing β-gal and Ptf1a expression in PWM at P1, P3, P6 and P8. Arrowheads indicate cells co-express β-gal and Ptf1a. Scale bars indicate 50 μm. (B and C) Quantification of total Ptf1a ^+^ cells (B) and the percentage of β-gal ^+^/Ptf1a^+^ double-positive cells (C) in the PWM. N= 4 mice per group. (D) Brains from *Smo* ^*F/+*^ and *Ptf1a*^*Cre/+;*^ *Smo*^*F/-*^ animals at P21 showing external views and H&E stained sections. (E and F) Quantification of the overall midsagittal area of the cerebellum and the molecular layer in P21 *Smo* ^*F/+*^and *Ptf1a*^*Cre/+;*^ *Smo* ^*F/-*^ mice. N= 3 to 5 mice per group. (G) Midsagittal cerebellar sections from P21 *Smo* ^*F/+*^and *Ptf1a*^*Cre/+;*^ *Smo* ^*F/-*^ mice stained with antibodies against parvalbumin (Parv). Scale bars indicate 50 μm. (H-J) Quantitative analysis of GABA^+^ DCN (H), Parv^+^ interneurons (I), Calbindin+ Purkinje cells (I) and Neurogranin+ Golgi cells (J) in P21 *Smo* ^*F/+*^and *Ptf1a*^*Cre/+;*^ *Smo* ^*F/-*^ cerebella. N= 3 to 5 mice per group. (K) The length of Purkinje cells’ dendritic spines in P21 *Smo* ^*F/+*^and *Ptf1a*^*Cre/+;*^ *Smo* ^*F/-*^ cerebella. N= 3 mice per group. All graphs displayed are mean ± SEM. *p ≤ 0.05, **p ≤ 0.01, ***p ≤ 0.001. n.s., not significant.

To ascertain the significance of Shh signaling in Ptf1a^+^ progenitors, we blocked Shh reception by genetically ablating the transducer Smo using the GABAergic lineage-specific *Ptf1a*^*cre*^ line (Kawaguchi *et al*, 2002). Our initial analysis of the mature cerebellar cortex of *Ptf1a*^*Cre*^; *Smo*^*F/-*^ mutant mice at P30 detected no structural or layering alterations; thickness of the molecular layer, granule layer, overall cerebellar area, and foliation pattern was comparable to the control (Figure 1D-1F). However, there is a clear reduction of hematoxylin/eosin-stained cells in the molecular layer. Indeed, immunohistochemical (IHC) detection of the calcium-binding protein, parvalbumin, which preferentially marks MLIs and PCs, showed a statistically significant decrease in the total number of MLIs in *Ptf1a*^*Cre*^; *Smo*^*F/-*^ mutant cerebella (Figure 1D and 1G). The most reduction was most prominent in the outer half of the ML, suggesting that a germinal program was biased towards the stellate interneuron subtype (see below). The number of other early-born inhibitory neurons, including deep cerebellar nuclei (DCN), PCs and Golgi cells expressing GABA, Calbindin and Neurogranin, respectively, was not significantly changed in *Ptf1a*^*Cre*^; *Smo*^*F/-*^ mutants (Figure 1H-1J). Additionally, no significant changes in Purkinje cell spine density were observed (Figure 1 K).

### Shh signaling regulates precursor pool expansion

To understand how Shh signaling affects the MLI pool size, we evaluated the proliferation and survival of Ptf1a progenitors in the PWM. Following a short-term (1h) labeling with the thymidine analog EdU, we observed a significant reduction of proliferating Ptf1a progenitors in *Ptf1a*^*Cre*^; *Smo*^*F/-*^ mutants at both P3 (∼20%) and P6 (∼34%) when compared to controls in the vermis (Figure 2A-2C). Accordingly, the total number of Ptf1a+ progenitors in the mutant was reduced by ∼30% and ∼68% at P3 and P6, respectively (Figure 2B and 2C). The proliferative effect of Shh signaling on Ptf1a+ progenitors was not restricted to the vermal region, as a similar reduction (∼39%) was also observed in parasagittal sections at P6 (Figure S1). Viability was measured with cleaved-Caspase-3, but no appreciable apoptotic cells were found in Ptf1a lineage cells at either P3 or P6 (Figure S2).

**Figure 2.**
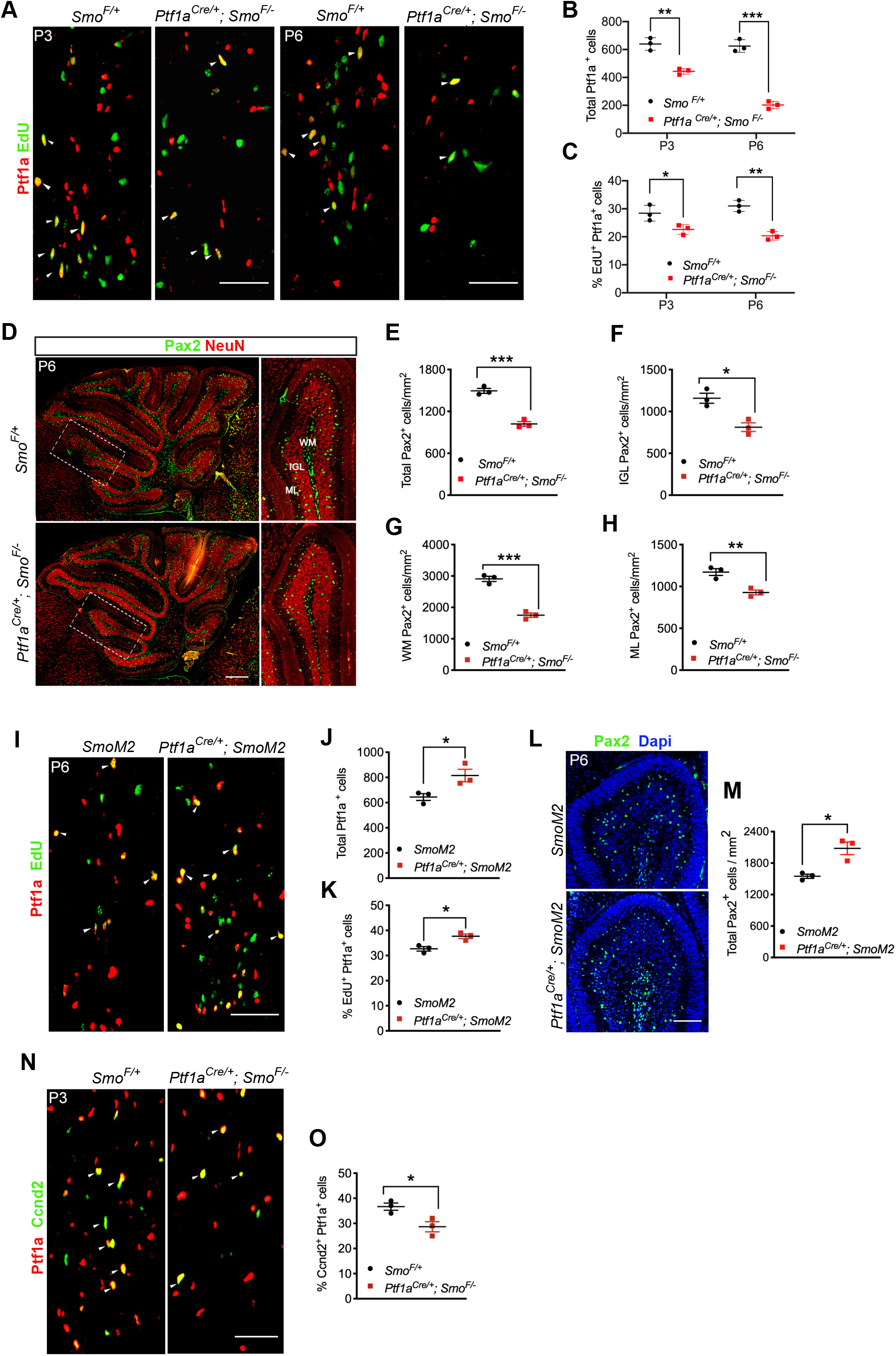
Shh signaling promotes Ptf1a+ progenitor proliferation and the expansion of Pax2^+^ immature interneurons. (A-C) Ptf1a and EdU staining in PWM of *Smo* ^*F/+*^and *Ptf1a*^*Cre/+;*^ *Smo* ^*F/-*^ cerebella at P3 and P6. Arrowheads indicate cells positive for both EdU and Ptf1a. Quantification of total Ptf1a ^+^ cells in the PWM at P3 and P6 (B). The percentage of EdU ^+^/Ptf1a^+^ double-positive cells relative to the total number of Ptf1a^+^ cells in PWM at P3 and P6 (C) is also quantified. N= 3 mice per group. Scale bars indicate 50 μm. (D) Pax2 and NeuN staining in sagittal sections of *Smo* ^*F/+*^and *Ptf1a*^*Cre/+;*^ *Smo* ^*F/-*^ cerebella. A higher magnification view of boxed regions is shown on the right. Abbreviations: WM, white matter; IGL, inner granule layer; ML, molecular layer. Scale bars indicate 100 μm. (E-H) Quantitative analysis of Pax2+ cells in different regions of the cerebellum shown in D. N= 3 mice per group. (I) Ptf1a and EdU staining in PWM of *SmoM2 and Ptf1a*^*Cre/+*^; *SmoM2* mice at P6. Arrowheads indicate cells positive for both EdU and Ptf1a. Scale bars indicate 50 μm. (J-K) Quantitative analysis of total Ptf1a ^+^ cells (J) and the percentage of EdU ^+^/Ptf1a^+^ double-positive cells relative to the total number of Ptf1a^+^ cells (K) in PWM at P6. (L) Pax2 staining in the ML of *SmoM2 and Ptf1a*^*Cre/+*^; *SmoM2* mice at P6. Scale bars indicate 100 μm. (M) Quantitative analysis of Pax2^+^ cells shown in L. n= 3 mice per group. (O) Ptf1a and Ccnd2 staining in PWM of *SmoM2 and Ptf1a*^*Cre/+*^; *Smo* ^*F/-*^ mice at P6. Arrowheads indicate cells co-expressing Ccnd2 and Ptf1a. Scale bars indicate 50 μm. (P) Quantitative analysis of the percentage of Ccnd2 and Ptf1a double-positive cells relative to the total number of Ptf1a^+^ cells in PWM at P6. N= 3 mice per group. All graphs displayed are mean ± SEM. *p ≤ 0.05, **p ≤ 0.01, ***p ≤ 0.001. n.s., not significant.

Genetic fate-mapping studies have shown that Ptf1a^+^ cells emerge upstream of and contribute considerably to neonatal Pax2^+^ immature interneuron pools (Fleming *et al*, 2013). The peak of Pax2 production in PWM occurs at P5, where they subsequently migrate to ML through the inner granule layer (IGL) before terminally differentiating into BCs and SCs (Weisheit *et al*, 2006; Wefers *et al*, 2018). We measured the abundance of Pax2^+^ cells at P6 and observed a significant reduction of their numbers in PWM, IGL, and ML (Figure 2D-H).

The role of Shh signaling in promoting Ptf1a progenitor proliferation was further evaluated using a gain-of-function approach. We generated *Ptf1a*^*cre*^; *SmoM2* mice by crossing the *Ptf1a*^*Cre*^ driver strain to *SmoM2* conditional mutants that harbor a constitutively activated form of *Smo* (Mao *et al*, 2006). Consistent with the loss of function study, we observed a significant increase in the number of Ptf1a+ progenitors (∼27%) as well as their proliferative capacity (∼15%) in *Ptf1a*^*cre*^; *SmoM2* cerebella when compared to controls at P6 (Figure 2I-2K). Accordingly, the number of Pax2+ cells is also significantly increased (Figure 2L and 2M). Together, we provide strong evidence for a direct and essential role of Shh signaling in driving the proliferation of Ptf1a+ progenitors in PWM and subsequent expansion of Pax2+ immature interneurons.

Previous studies have shown that *Cyclin d2* (*Ccnd2*) is expressed in the PWM and required for the proliferation of MLI progenitors (Huard *et al*, 1999), but the signal that activates its expression is unknown. We therefore examined Ccnd2 expression and found that the number of Ccnd2+ GABAergic progenitors is significantly reduced in the *Ptf1a*^*Cre*^; *Smo*^*F/-*^ PWM by immunohistochemistry (Figures. 2O and 2P). These data suggest that Shh signaling regulates *Ccnd2* expression that contributes to proliferation of Ptf1a progenitors.

### Specification of MLI subtypes occurs independent of their birth orders

To precisely determine whether specific MLI subtypes are affected in *Ptf1a*^*Cre*^; *Smo*^*F/-*^ mutants, we acquired the *Ret*^*GFP*^ mouse line in which EGFP reporter is expressed from the endogenous Ret promoter (Jain *et al*, 2006). Consistent with previous studies (Sergaki *et al*, 2017), *Ret*^*GFP*^ expression in BCs is first detectable at P7 and continues throughout adulthood (Figure S3A). Outside of the ML, *Ret*^*GFP*^ is expressed in the deep cerebellar nuclei (DCN) and associated fibers in PWM (Figure S3B and S3C), but not in Pax2+ interneuron progenitors as reported previously (Sergaki *et al*, 2017). The Ret+ DCN neurons are NeuN positive but GABA negative, indicating that they are not GABAergic neurons (Figure S3D). Therefore, *Ret*^*GFP*^ expression is confined to mature BCs and DCN neurons. In addition to *Ret*^*GFP*^ expression, we incorporated Parvalbumin (Parv) as a marker to highlight all MLIs. In this way, BCs and SCs can be unambiguously identified as Ret^GFP+^ Parv^+^ and Ret^GFP−^ Parv^+^ cells, respectively. Using these markers, we were able to determine that SC numbers were reduced by ∼21% in *Ptf1a*^*Cre*^; *Smo*^*F/-*^ mutant cerebella (Figure 3A and 3B) whereas no appreciable changes in BC numbers were observed when compared to that of controls (Figure 3A and 3C).

**Figure 3.**
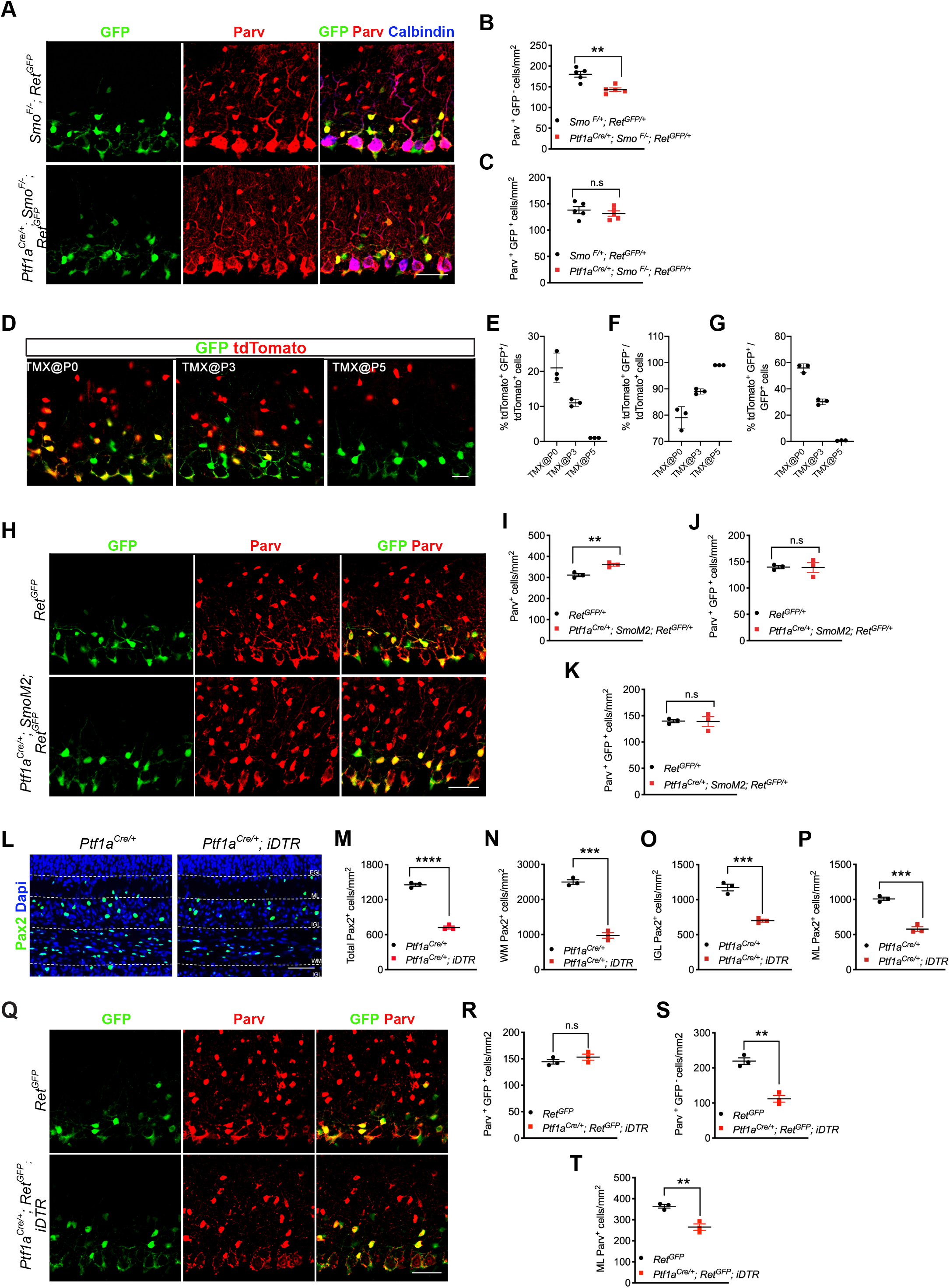
Shh signaling selectively regulates stellate cell pool size. (A) Cerebellar sections from P21 *Smo* ^*F/+*^; *Ret*^*GFP/+*^ *and Ptf1a*^*Cre/+*^; *Smo*^*F/-*^; *Ret*^*GFP/+*^ mice stained with antibodies against GFP, Parvalbumin and Calbindin. (B-C) Quantitative analysis of Parv^+^ GFP^+^ (B) and Parv^+^ GFP^−^ (C) cells shown in A. N= 5 mice per group. (D) Cerebellar sections from P21 *Ptf1a*^*Cre/+*^; *R26R*^*Ai9*^; *Ret*^*GFP/+*^ mice showing the extent of BC labeling (dtTomato^+^ GFP^+^) following TMX induction at P0, P3 or P5. (E-F) Quantitative analysis of the percentage of labeled BCs (dtTomato^+^ GFP^+^) and SCs (dtTomato^+^ GFP^−^) relative to total labeled (dtTomato^+^) cells. (G) Quantitative analysis of the percentage of labeled BCs (dtTomato^+^ GFP^+^) relative to total GFP+ BCs. (H) Cerebellar sections from P21 *Ret*^*GFP/+*^ *and Ptf1a*^*Cre/+*^; *SmoM2; Ret*^*GFP/+*^ mice stained with antibodies against GFP and Parvalbumin. (I-K) Quantitative analysis of Parv+ (I), Parv^+^ GFP^+^ (J) and Parv^+^ GFP^−^ (K) cells shown in H. N= 3 mice per group. (L) Cerebellar sections from P6 *Ret*^*GFP/+*^ and *Ptf1a*^*Cre/+*^; *iDTR* mice showing Pax2 distribution (green) in EGL, IGL and WM following DT administration at P0-P2. (M-P) Quantitative analysis of total Pax2+ cells (M) and Pax2+ cells in WM (N), IGL (O) and ML (P). N= 3 mice per group. (Q) Cerebellar sections from P21 *Ret*^*GFP/+*^ and *Ptf1a*^*Cre/+*^; *iDTR* mice stained with antibodies against GFP and Parvalbumin. (R-T) Quantitative analysis of Parv^+^ GFP^+^ (R), Parv^+^ GFP^−^ (S) and Parv^+^ (T) cells shown in A. N= 3 mice per group. All graphs displayed are mean ± SEM. *p ≤ 0.05, **p ≤ 0.01, ***p ≤ 0.001. n.s., not significant.

Classical birthdating and genetically inducible fate mapping (GIFM) studies have suggested that MLI subtype identities are correlated with their birth dates and laminar positions within the cerebellar cortex (Leto *et al*, 2006; Sudarov *et al*, 2011). To determine if the subtype-specific effect of Shh signaling on MLIs is associated with temporal differences in Shh responsiveness of Ptf1a^+^ progenitors fated to generate both BCs and SCs, we generated *Ptf1a*^*creER*^; *Ai9; Ret*^*GFP*^ line to perform GIFM study focusing on critical stages where Shh signaling is activated in Ptf1a progenitors. Tamoxifen (TM) administration at P0 marked both BCs (dtTomato^+^ Ret^GFP+^) and SCs (dtTomato^+^ Ret^GFP−^) in the molecular layer at P21, representing ∼21% and ∼79% of all marked cells, respectively (Figure 3D, 3E and 3F). Moreover, marked BCs account for ∼56% of all Ret^GFP+^ cells (Figure 3G), indicating that the peak of BC production occurs between P0 and P1. When TM was administered at P3, marked BCs decreased to ∼30% (Figure 3G), with a concomitant increase in marked SCs (Figure 3F). At P5, SCs are the only labeled population present in the ML (Figure 3F and 3G). Therefore, BCs and SCs are both generated at the time when Shh signaling is active in Ptf1a+ progenitors, consistent with previous *Gli1*^*CreER*^ GIFM studies in which BC and SC subtypes are both generated from Shh-responsive progenitors (Fleming *et al*, 2013). Therefore, the differential effect of Shh signaling on MLI subtypes is unlikely due to temporal or regional difference in Shh responsiveness. Indeed, analysis of *Ptf1a*^*cre*^; *SmoM2* mutants with constitutive Shh pathway activation in Ptf1a^+^ progenitors showed that only Ret^GFP−^ Parv^+^ (SC) numbers are elevated whereas the Ret^GFP+^ Parv^+^ (BC) pool remained unchanged when compared to the control (Figure 3H-K).

The observation that altered numbers of Pax2^+^ immature interneurons in the gain and loss of Shh signaling mutants had no effect on BC pool size suggests commitment to the BC fate from Pax2+ cells does not occur until they settle into the inner ML. Accordingly, the BC pool remains unchanged as long as there are sufficient Pax2^+^ cells to fill the inner ML. To further test this model, we generated *Ptf1a*^*cre*^; *Ret*^*GFP*^; *ROSA26*^*DTR*^ mice designed to temporarily ablate Ptf1a^+^ progenitors and their descendent Pax2^+^ cells. The *ROSA26*^*DTR*^ line expresses the Cre-inducible Diptheria toxin receptor (DTR) that triggers cell ablation upon exposure to Diptheria toxin (DT). Administration of DT at P0-P2, a time when Ptf1a^+^ progenitors are fated to generate most BCs, resulted in a drastic reduction of Pax2^+^ immature interneurons in all layers of the cerebellum at P6 (Figure 3L-3P). Despite ∼50% reduction of total Pax2^+^ cells (Figure 3M), we observed no effect on BC numbers while SCs are reduced by ∼49% at P21 (Figure 3Q – 3T). The lack of perturbations in BC numbers is not due to selective depletion of Ptf1a^+^ progenitors at the later time period when SCs are generated as evidenced by more than 27% reduction of Ptf1a+ progenitors at P2 when DT was administered at P0 and P1 (Figure S4). We also ruled out the possibility of a delay in BCs to SCs production in DT treated mice, which would have prolonged BCs production and preferentially affected SC numbers. For this, we performed classical birthdating study on DT treated pups (P0-P2) by administrating EdU at P5 when BCs are no longer generated (Figure 3D-G). Analysis of EdU labeled cells at P21 revealed that only SCs (Ret^−^ Parv^+^) but no BCs (Ret^+^ Parv^+^) were labeled after DT treatment (Figure S4), indicating that the progenitor cell ablation did not elicit a delay in BCs to SCs production. Collectively, the results suggest that MLI subtypes are specified independent of their birth order but likely occur at their laminar positions, providing a mechanism by which Shh signaling is selectively promoting the expansion of SCs.

### Impaired GABAergic synapses and inhibitory control over PCs

In the mature cerebellum, SCs and BCs relay inhibitory input to PCs via direct synapses on dendritic tree or somata/proximal initial segment, respectively, balancing excitatory input received from granule neurons and afferent fibers (Sotelo, 2015). To understand how the loss of Shh-dependent SCs might impact the integrity of cerebellar neural circuitry, we evaluated the distribution of specific molecular markers for inhibitory and excitatory synapses in the ML. We used a presynaptic marker, vesicular GABAergic transporters (VGAT), in combination with a postsynaptic marker Gephyrin to highlight inhibitory synapses. In the ML, the number of inhibitory synapses (puncta with co-localization) at PC dendrites was reduced by 31% in *Ptf1a*^*Cre*^; *Smo*^*F/-*^ mutants compared to controls (Figures 4A-4C). We also used another inhibitory presynaptic marker, glutamic acid decarboxylase 67 (Gad67), which is responsible for up to 90% of GABA synthesis in the brain (Asada *et al*, 1996; Kash *et al*, 1997), and found that its puncta are also significantly reduced (Figure S4). Consistent with unperturbed BC development, the number of inhibitory synapses at PC soma and the pinceau marked by the potassium voltage-gated channel 1.2 (Kv1.2) remains similar between *Ptf1a*^*Cre*^; *Smo*^*F/-*^ mutants and controls (Figures 4D and 4E). However, VGluT1 (Figures 4F and 4H), which labels excitatory synaptic terminals at granule cell parallel fibers, showed no significant changes, nor did VGluT2 (Figures 4G and 4I), which marks excitatory synaptic terminals at climbing fibers. Altogether these findings show substantial impairment to the inhibitory component of the cerebellar system in *Ptf1a*^*Cre*^; *Smo*^*F/-*^ mutants, while the excitatory component appears to be unperturbed.

**Figure 4.**
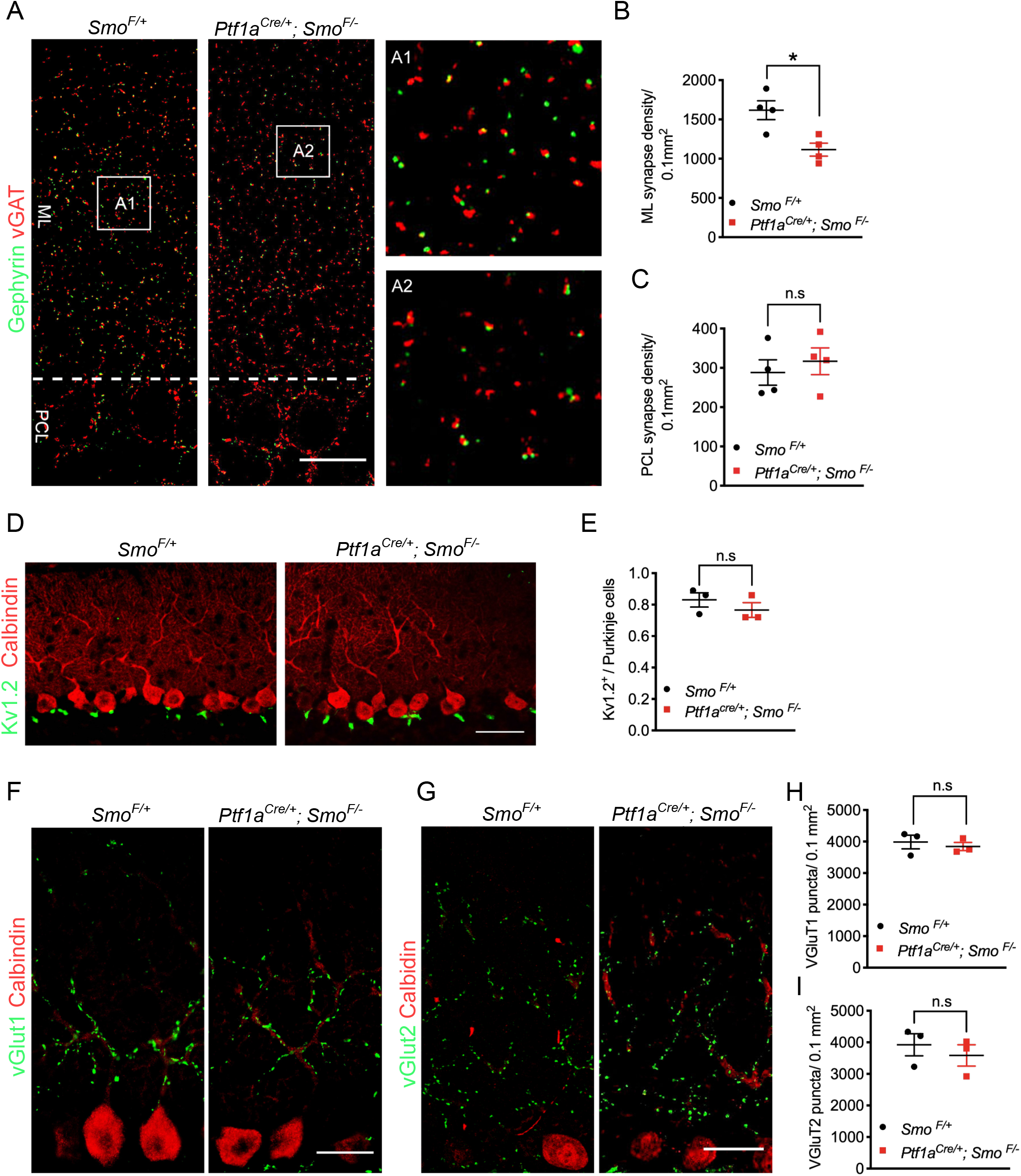
Loss of Shh-dependent stellate cells reduces ML GABAergic synapses. (A-A2) Cerebellar sections from P21 *Smo* ^*F/+*^ *and Ptf1a*^*Cre/+*^; *Smo*^*F/-*^ mice stained with antibodies against vGAT and Gephyrin. A1 and A2 represent enlarged views of the boxed regions in A. ML, molecular layer. PCL, Purkinje cell layer. Scale bars indicate 25 μm. (B-C) Quantitative analysis of vGAT and Gephyrin double-positive synapses in the ML (B) and PCL (C). N= 4 mice per group. (D) Cerebellar sections from P21 *Smo* ^*F/+*^ *and Ptf1a*^*Cre/+*^; *Smo*^*F/-*^ mice stained with antibodies against Kv1.2 and calbindin. Scale bars indicate 50 μm. (E) Quantitative analysis of Kv1.2 in PCL shown in D. N= 4 mice per group. (F-G) Cerebellar sections from P21 *Smo* ^*F/+*^ *and Ptf1a*^*Cre/+;*^ *Smo*^*F/-*^ mice stained with antibodies against vGluT1 (F), vGluT2 (G) and calbindin. Scale bars indicate 25 μm. (H and I) Quantitative analysis of vGluT1 and vGluT2 positive puncta in the ML. N= 3 mice per group. All graphs displayed are mean ± SEM. *p ≤ 0.05, **p ≤ 0.01, ***p ≤ 0.001. n.s., not significant.

The significantly reduced number of dendritic inhibitory synapses in *Ptf1a*^*Cre*^; *Smo*^*F/-*^ mice suggests impaired neurotransmission. To directly test this, we performed whole-cell patch-clamp recordings on cerebellar slices to compare spontaneous inhibitory postsynaptic currents (sIPSCs) of PCs as a measure of inhibitory synaptic strength in wild-type (n=10) and mutant (n=8) mice. This analysis revealed that adult *Ptf1a*^*Cre*^; *Smo*^*F/-*^ mice had about a 35% decrease in the frequency of sIPSCs when compared to the control (Figure 5A-5C). Similarly, the amplitude of sIPSCs is also significant reduced in *Ptf1a*^*Cre*^; *Smo*^*F/-*^ mice (Figure 5B). These findings indicate that PCs lose dendritic synapses and show attenuated inhibitory input.

**Figure 5.**
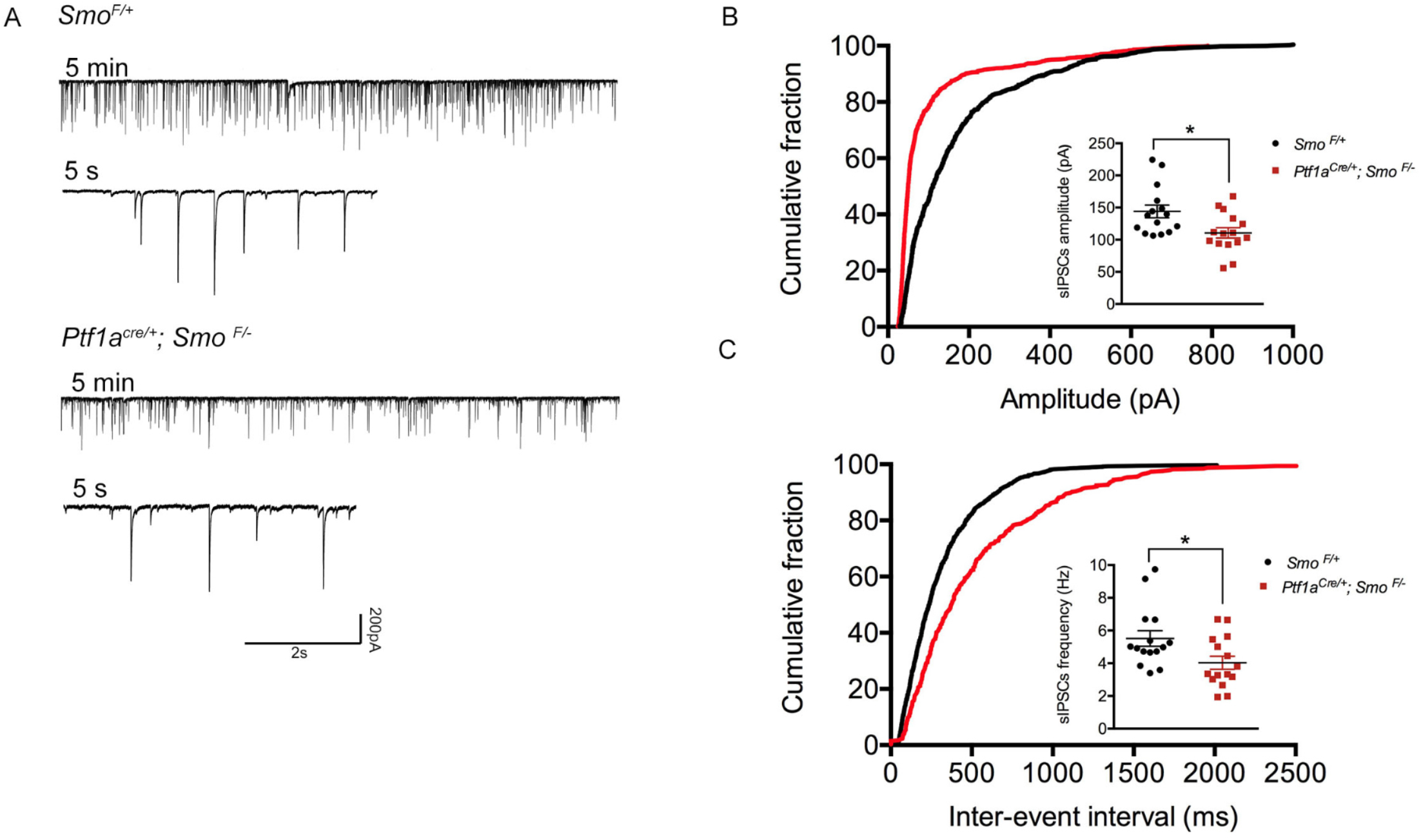
Decreased amplitude and interevent interval of sIPSCs in *Ptf1a*^*Cre/+*^; *Smo*^*F/-*^mutants. (A) Representative traces of spontaneous IPSC recorded from *Smo* ^*F/+*^ *and Ptf1a*^*Cre/+*^; *Smo* ^*F/-*^ cerebellar Purkinje cells. (B-C) Statistical analysis of sIPSCs amplitude (B) interevent interval (C). Each dot represents the mean of values obtained from 5 min recording sessions of individual PC. N= 15 cells from 11 *Smo* ^*F/+*^ mice and 8 *Ptf1a*^*Cre/+*^; *Smo* ^*F/-*^ mice. Values represent mean ± SEM. * p < 0.05.

### Diminished capacity for associative motor learning

To determine whether the reduction of dendritic inhibitory neurotransmission on PCs might impair cerebellum-dependent functions, a series of neurobehavioral assays were performed. The cerebellum is classically known to coordinate motor function, and the most obvious manifestation of cerebellar impairment is abnormal gait. *Ptf1a*^*Cre*^; *Smo*^*F/-*^ mice did not demonstrate gross impairments in mobility, such as ataxia, nor were they observed to show clinical signs of tremor. When subjected to treadscan analysis, adult *Ptf1a*^*Cre*^; *Smo*^*F/-*^ (n=12) and control mice (n=11) showed comparable gait, which is divided into three basic components: i) stance (break + propulsion), ii) swing, and iii) stride (stance + swing), and is measured in both length and time (Figures 6A-6E). To test motor learning, these animals were also subjected to an accelerating rotarod assay. On the first day of testing sessions, mutant and control animals showed nearly identical latency to fall, indicating that no obvious differences in motor coordination, balance, or strength were detectable (Figure 6F). However, during the subsequent days, it appeared that *Ptf1a*^*Cre*^; *Smo*^*F/-*^ mutants had an apparent deficiency in the ability to learn to perform this task, with statistically significant shorter latencies to fall than controls (Figure 6F). Overall, loss of a SC subset did not appear to adversely impact gross motor function in *Ptf1a*^*Cre*^; *Smo*^*F/-*^ mutants and only imparted a subtle impairment to primitive motor learning.

**Figure 6.**
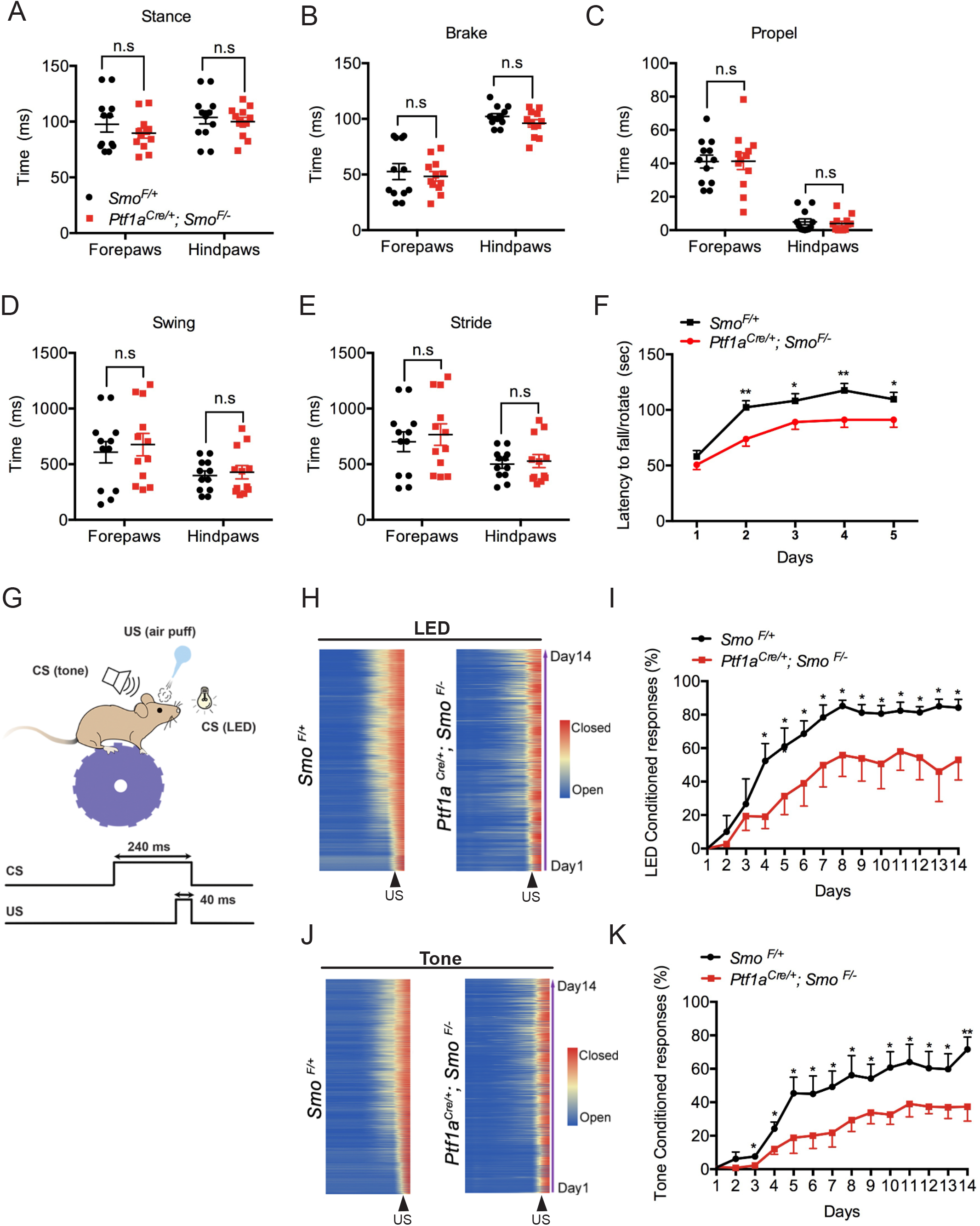
Shh-dependent SC pool is required for cerebellum-dependent motor learning but not basic motor function. (A-E) Quantitative gait analysis of adult *Smo* ^*F/+*^ and *Ptf1a*^*Cre/+*^; *Smo*^*F/-*^ mice showing stance time (A), brake time (B), propel time (C), swing time (D) and stride time (E). N= 12 mice per group. (F) Accelerating rotarod test to measure latency to fall on five consecutive days in *Smo* ^*F/+*^ and *Ptf1a*^*Cre/+*^; *Smo*^*F/-*^ mice. N= 30 mice per group. (G) Schematic of eyeblink conditioning system. The trial structure is consists of a 240 ms blue light (CS) or tone (CS) that precedes and co-terminates with a 40 ms air puff (US). (H-J) Group averages of trial-by-trial eyelid closure over the entire 14 days of conditioning with LED (H) or tone (J) in *Smo* ^*F/+*^ and *Ptf1a*^*Cre/+*^; *Smo*^*F/-*^ mice. (I-K) The percentage of conditioned responses LED (I) or tone (K). N= 5 mice per group. All graphs displayed are mean ± SEM. * p < 0.05, **p < 0.01. n.s., not significant.

Proper cerebellar function is also necessary for delay eyeblink conditioning, a form of associative motor learning required for multi-sensory integration that is often impaired in autistic individuals (Medina *et al*, 2000; Sears *et al*, 1994). The paradigm involves pairing a neutral conditioned stimulus (CS; e.g., LED light pulse or tone) with an eyeblink-evoking unconditioned stimulus (US; e.g., an air puff). After repeated CS-US pairing, an association is progressively established such that the eyelid conditioned response (CR), in the form of learned blink, occurs before the onset of the US. We, therefore, determined whether cerebellum-dependent eyeblink conditioning is compromised in *Ptf1a*^*Cre*^; *Smo*^*F/-*^ mice with reduced SC numbers. In this paradigm, a blue LED light pulse was used as the CS for a period of 240 ms followed immediately by a gentle air puff for 40 ms (Figure 6G). Eyelid movement was measured using a high-speed infrared camera interfaced with MATLAB (Heiney *et al*, 2014) and the eyelid position was assigned to a value between 0 (fully open) and 1 (fully closed). After 14 days of consecutive sessions of conditioning (1400 trials in total), *Ptf1a*^*Cre*^; *Smo*^*F/-*^ mice had a consistently lower amplitude of eyelid closure just before CS exposure when compared to control mice (Figure 6H and 5S). Although both the mutant and control mice showed a gradual increase in the percentage of CRs, mutant mice had significantly lower CRs throughout the conditioning sessions (Figure 6I).

While the Shh pathway has not been reported to be active in the retina of Ptf1a-expressing cells and their lineage, we were concerned that the CR deficit in *Ptf1a*^*Cre*^; *Smo*^*F/-*^ mutants is due to compromised retina function. We, therefore, performed electroretinograms (ERG) and visual evoked potentials (VEP) to assess retinal and visual pathway activity in response to light. We observed no significant differences in maximum responses of rod photoreceptors (a_max_), inner retinal neurons (b_max_ and oscillatory potentials; OP), or the visual cortex (VEP N1) between *Ptf1a*^*Cre*^; *Smo*^*F/-*^ and control mice (Figure S6), indicative of normal retinal and visual pathway function. To confirm that the mutants have a deficit in associative motor learning, we also used a tone as CS in delayed eyeblink conditioning. Similar to LED light, the mutant mice showed a diminished response to the tone as presented by the lower amplitude of eyelid closure and percentage of CRs (Figure 6J, 6K, S5). Collectively, these results indicate that Shh signaling-dependent dendritic synapses formation plays a critical role in associative motor learning.

## DISCUSSION

MLIs constitute the majority of inhibitory interneurons in the cerebellum, but the mechanisms that regulate their subtype identities and pool sizes remain not well understood. We reveal that Shh signaling is activated in a subset of interneuron progenitors that give rise to both BCs and SCs, but surprisingly it is selectively required for the expansion of the SC pool. Through lineage tracing, genetic gain and loss of function, and cell ablation experiments, we show that molecular layer interneuron subtypes are specified independent of Shh signaling and their birth orders but appear to occur in their terminal laminar positions according to inside-out sequence. Our studies at the synaptic and behavioral levels show that dendritic GABAergic inhibition controlled by Shh signaling-dependent SC pools is critical for motor learning.

Previous studies have shown that MLIs originate from multipotent astroglia in the PWM (Fleming *et al*, 2013; Parmigiani *et al*, 2015; Silbereis *et al*, 2009). These astroglia act as resident stem-like cells and proliferate in response to the Shh signal from PCs to generate astrocyte precursors as well as Ptf1a progenitors (Fleming *et al*, 2013). Our finding that Shh signaling is directly required for the proliferation of Ptf1a+ progenitors and subsequent expansion of MLIs offers an additional mechanism by which Shh regulates MLI pool size.

It is estimated that approximately 90% of all inhibitory neurons in the cerebellum are represented by late-born GABAergic interneurons (Weisheit *et al*, 2006). These neurons come from Pax2-expressing immature interneurons that are mostly generated during the first week of postnatal life (Weisheit *et al*, 2006). As Pax2^+^ cells descend from Ptf1a^+^ progenitors (Fleming *et al*, 2013) and emerge at the final progenitor cell division, Shh-induced proliferation of Ptf1a+ progenitors, therefore, serves as the main driver for their rapid expansion. This mode of regulation is in contrast to other regions of the nervous system where Shh pathway activity is generally excluded from postmitotic Ptf1a expressing cells. In the spinal cord, Ptf1a expression is restricted to the postmitotic progenitors in the dorsal neural tube whereas the Shh pathway is activated in the ventral domain (Glasgow *et al*, 2005; Bai *et al*, 2004). Moreover, ectopic Ptf1a expression is correlated with the downregulation of the Shh signaling pathway as observed in Danforth’s short tail mice (Orchard *et al*, 2018). Similarly, in the developing retina, Shh signaling is activated in early retinal progenitors whereas Ptf1a expression is restricted to postmitotic horizontal and amacrine cells (Fujitani *et al*, 2006). Indeed, the lack of Shh pathway activity in postmitotic retinal lineage cells is consistent with the absence of visual impairment in *Ptf1a*^*cre/+*^; *Smo*^*F/-*^ mice (Figure S5). Therefore, the Shh signaling pathway appears to be co-opted for the rapid expansion of Ptf1a^+^ progenitors to ensure an adequate supply of MLIs. Mechanistically, Ccnd2 is an attractive target that mediates the proliferative effect of Shh pathway activation on MLI progenitors. Unlike Ccnd1, Ccnd2 is strongly expressed in MLI progenitors and loss of Ccnd2 impairs their production (Huard *et al*, 1999). Reduced expression of Ccnd2 in Smo-deficient Ptf1a progenitors is consistent with the Shh pathway regulating Ccnd2 expression.

It has been reported that cerebellar MLIs undergo limited programmed cell death during the first two weeks of postnatal development (Yamanaka *et al*, 2004). More recent studies showed that GDNF signaling is required for the survival of MLIs. Genetic deletion of GDNF receptor GFRa1 or Ret in GABAergic progenitors resulted in a ∼25% reduction of MLIs (Sergaki *et al*, 2017). Interestingly, the loss of NeuroD2, a bHLH transcription factor required for MLI differentiation, also promotes MLI survival (Pieper *et al*, 2019). Thus, MLI pool size is regulated at multiple levels by distinct mechanisms, from the proliferation of multipotent astroglia and Ptf1a progenitors to the survival of MLIs.

Fate mapping and heterochronic transplantation studies have suggested that MLI identities and laminar placement link to birthdate within the cerebellar cortex (Altman & Bayer, 1997; Cameron *et al*, 2009; Leto *et al*, 2009; Sudarov *et al*, 2011). Accordingly, a reduction in Ptf1a+ progenitors and Pax2+ immature interneurons as observed in *Ptf1a*^*cre/+*^; *Smo*^*F/-*^ mutants would have a profound effect on the production of both BCs and SCs. It is, therefore, unexpected to discover that *Ptf1a*^*cre/+*^; *Smo*^*F/-*^ mutants did not affect BC production. This is not due to a lack of Shh responsiveness of Ptf1a+ progenitors fated to generate BCs as our *Ptf1a*^*CreER*^ GIFM study showed that BCs are largely generated from P0 to P3 when the majority of Shh responsive Ptf1a+ progenitors are present. Importantly, *Gli1*^*CreER*^ GIFM studies have shown that both MLI subtypes are generated from Shh-responsive progenitors (Fleming *et al*, 2013). The fact that the increased Pax2+ cell number observed in *Ptf1a*^*cre/+*^*;SmoM2* mutants did not affect BC pool size further suggests that the commitment to BC and SC fates is not directly related to their birth order but appears to occur at laminar positions in an inside out sequence. Accordingly, Pax2^+^ cells that initially populate the inner ML acquire BC identity whereas those remaining in the outer ML assume SC fate. This model is further supported by our genetic ablation experiment in which the reduction of Ptf1a+ progenitors at the peak of BC production did not affect BC numbers despite a drastically diminished number of Pax2^+^ cells. We have ruled out the possibilities that the lack of BC perturbation is due to a delay in the progenitor cell depletion or timing of BC to SC production. Thus, changes in Pax2^+^ pool sizes as observed in Smo gain and loss-of-function mutants will have the greatest impact on SC numbers. Our model is consistent with the plastic nature of cerebellar immature interneurons observed from heterochronic transplantation studies(Leto *et al*, 2009), in which P7 interneurons grafted to P1 cerebella exclusively adopt basket cell fate. Our model also explains why the preferential reduction of SCs in the ML is observed in *Ccnd2* and *Ascl1* mutants despite an early loss of interneuron progenitors (Huard *et al*, 1999; Sudarov *et al*, 2011).

Recent studies have shown that immature GCs play an instructive role in differentiation of SCs at the ML (Cadilhac *et al*, 2021). At present, Ret is the only marker that is selectively expressed in the BCs. Thus, understanding how its expression is activated may provide insight into how laminar positional information affects BC’s identity. Ret has been studied extensively in the context of kidney and enteric nervous system development (Lake & Heuckeroth, 2013; Costantini, 2016). Its expression in the ureteric bud and enteric neural crest-derived precursor cells is activated by retinoic acid (RA) signaling (Mendelsohn *et al*, 1999; Simkin *et al*, 2013). However, analysis of RA signaling using RA responsive reporter mice indicated that it is only activated in a subset of PCs, not in the BCs (Figure S7). Another possibility is that BCs identity is specified through an activity-dependent mechanism as proposed for the cortical interneuron specification (Wamsley & Fishell, 2017). There is evidence that the patterning of BC pinceau is shaped by PC activity (Zhou *et al*, 2020). In this context, Ret expression may be activated when the pinceau is established between its axon collaterals and the PC axon initial segment (AIS). Future studies are required to determine how positional information influences BC identity.

Previous studies have shown that the reduction of MLIs or the loss of their postsynaptic receptors γ2 GABA-A resulted in an impaired motor learning (Sergaki *et al*, 2017; Wulff *et al*, 2009; Brinke *et al*, 2015). However, it is unclear to what extent each MLI subtype contributes to motor learning. There are critical anatomical and functional differences in how SCs and BCs provide feed-forward inhibition of PCs in response to PF and CF activation. SCs primarily transmit chemical inhibition through axo-dendritic synapses whereas BCs deliver both chemical and electrical field inhibition through axo-somatic and axo-axonic synapses, respectively (Palay & Chan-Palay, 1974b; Korn & Axelrad, 1980; Blot & Barbour, 2014). Recent tamoxifen-mediated temporal deletions of the GABAergic transporter have suggested that BCs and SCs may have a distinct function in regulating PC simple spike firing pattern and rate (Brown *et al*, 2019). Our analysis of Shh signaling mutants provides evidence that SCs contribute significantly to motor learning. We found that the 21% reduction of SC numbers had a profound consequence on the cerebellar circuitry, resulting in a nearly 31% reduction in dendritic inhibitory synapses and significant impairment of inhibitory neurotransmission. This impairment in dendritic synapses and GABAergic input onto PCs is sufficient to cause motor learning deficit. It remains to be determined whether axo-somatic and axo-axonic synapses contribute to motor learning.

## ACKNOWLEDGEMENTS

We thank Jingqiong Kang, Ph.D. and Chun-Qing Zhang, M.D. for their initial help with electrophysiology experiments. We thank Christopher Wright, D.Phil. for the generous gift of antibody against Ptf1a. This study was supported by grants to C.C. from the Vanderbilt-Ingram Cancer Center Support Grant P30 CA068485 and the National Institutes of Health NS 097898.

## AUTHOR CONTRIBUTIONS

C.C. designed the experiments and wrote the paper with the help of W.L. and L.C. W.L. performed all the experiments except eyeblink conditioning and ERG. W. L. performed the electrophysiology experiment with the help of K.A.Z. and A.H.L. L.C., W.L. and J.T.F. performed eyeblink conditioning experiment with the help of S.A.H., G.J.W. and J.F.M. The ERG data was provided by T.S.R. C.C., W.L. and L.C. analyzed data.

## DECLARATION OF INTERESTS

The authors declare no competing interests.

## KEY RESOURCES TABLE

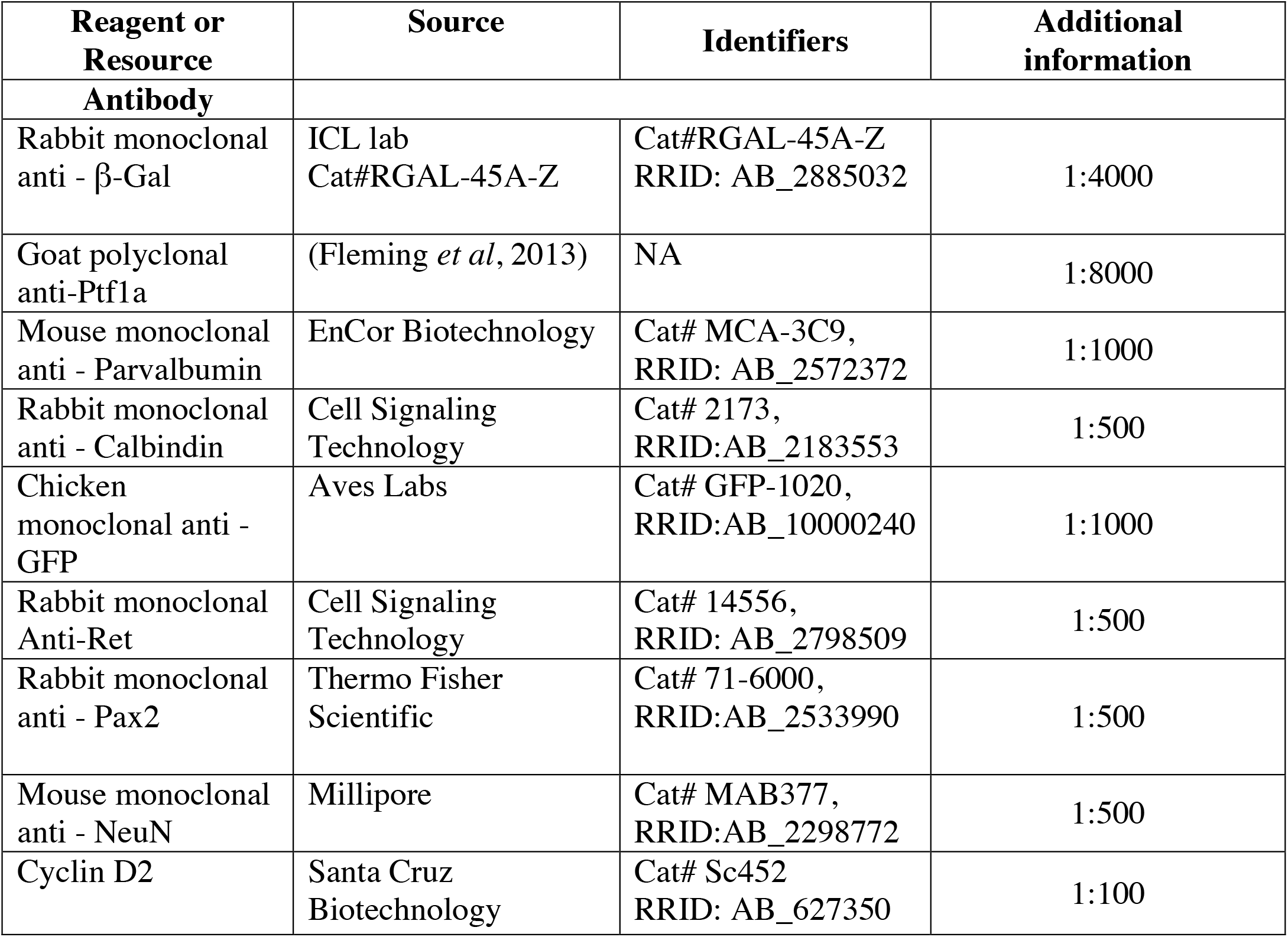
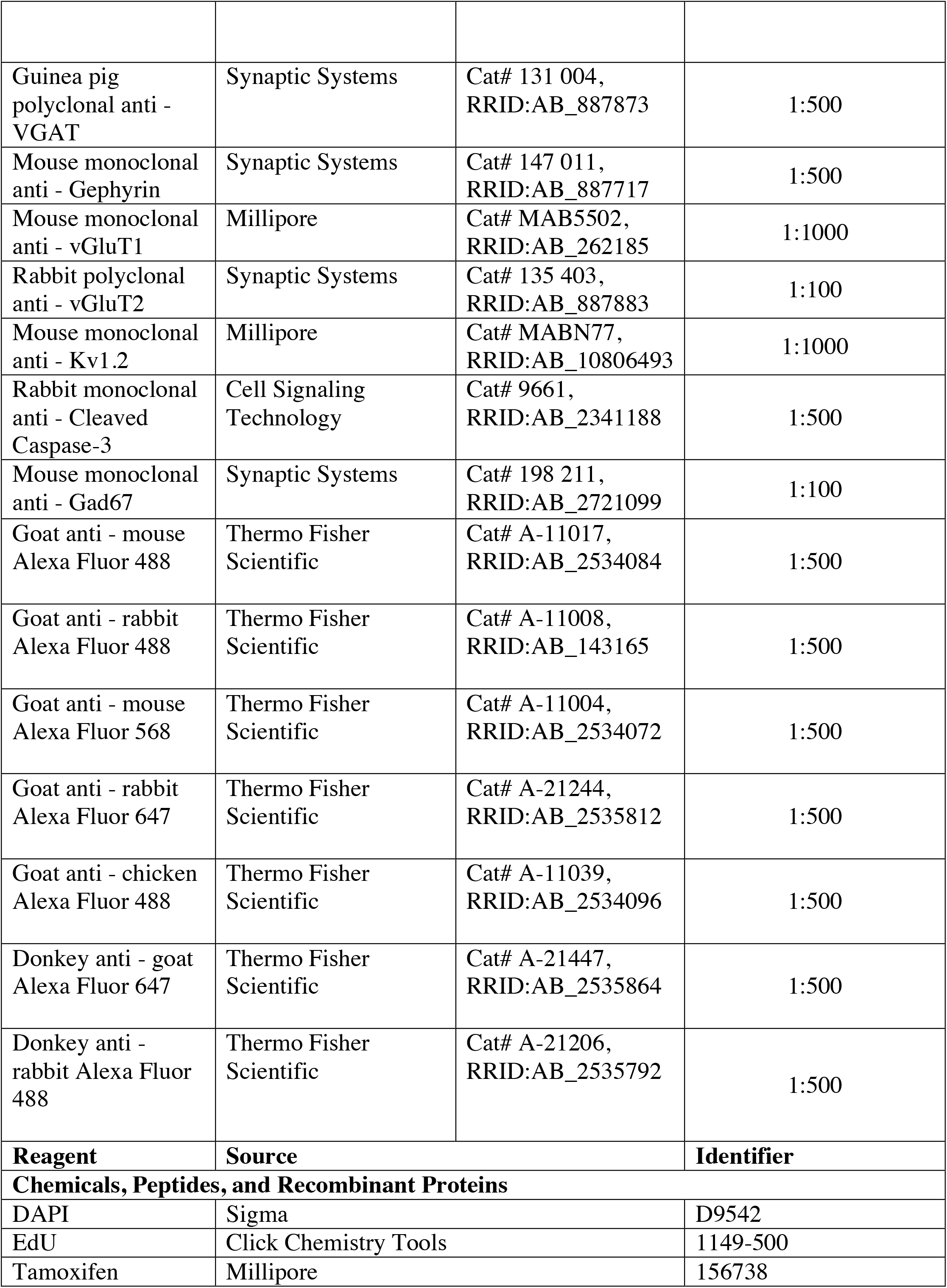
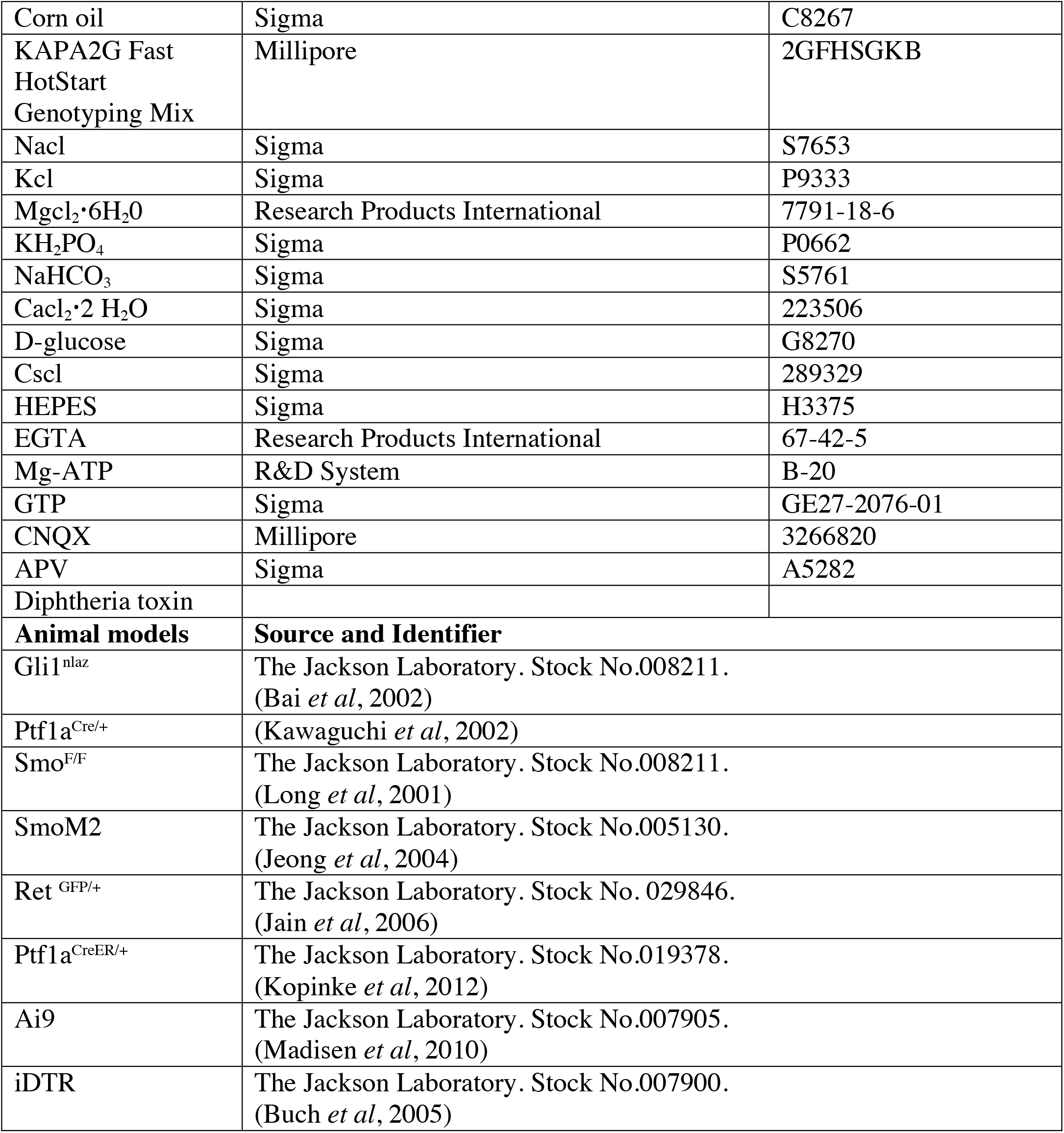

## EXPERIMENTAL PROCEDURES

### Mice

All procedures followed animal care and biosafety protocols approved by Vanderbilt University Division of Animal Care in accordance with NIH guidelines. Histological analyses were carried out at postnatal (P) 1 to P30. Behavior experiments used 3 to 5 months adult mice. The following mouse lines were used in this study:

1. *Gli1*^*nlaz*^, nlacZ reporter was inserted into the endogenous *Gli1* locus, creating a null allele and activating nlacZ expression in a pattern indistinguishable from that of *Gli1* (Bai *et al*, 2002). 2. *Ptf1a*^*Cre/+*^, Cre was inserted into the endogenous Ptf1a locus, creating a null allele and activating Cre expression in a pattern indistinguishable from that of *Ptf1a* (Kawaguchi *et al*, 2002). 3. *Ptf1a*^*CreER/+*^, CreER was inserted into the endogenous Ptf1a locus, creating a null allele and activating CreER expression in a pattern indistinguishable from that of *Ptf1a* (Kopinke *et al*, 2012). 4. *Smo*^*F/F*^, the loxP sequence was inserted on either side of exon 1 at the Smo locus, permitting a Cre recombinase dependent *Smo* loss of function mice (Long *et al*, 2001). 5. *R26*^*SmoM2*^, *a lox-stop-lox eYFP SmoM2* cassette was inserted into *ROSA26* locus, permitting the expression of eYFPSmoM2 fusion protein in a Cre dependent manner (Jeong *et al*, 2004). 6. *Ret* ^*GFP/+*^, exon 1 of the *Ret* locus was replaced by eGFP, creating a null allele and activating eGFP expression in a pattern similar to that of *Ret* (Jain *et al*, 2006). 7. *R26*^*iDTR*^, a loxp cassette containing diphtheria toxin receptor (DTR) was inserted into *ROSA26* locus, permitting the expression of DTR in a Cre dependent manner (Buch *et al*, 2005). 8. *R26R*^*Ai9*^, a *CAG lox-stop-lox tdtomato* cassette was inserted into *ROSA26* locus, permitting the expression of tdtomato under the control of CAG promoter in a Cre-dependent manner (Madisen *et al*, 2010).

### Mice treatments

For proliferation analysis, P3 and P6 pups were injected intraperitoneally with 50 mg/kg of EdU (5-ethyl-2-deoxyuridine; Sigma). Brains were harvested 2h after injection and processed for frozen tissue sections. Fate mapping studies, P0, P3 and P5 pups were injected intraperitoneally with 50 mg/kg of EdU (P5 pups only) or 50 mg/kg of tamoxifen (Sigma) dissolved in corn oil. Brains were harvested at P21 and processed for frozen tissue sections as described below. P0 Pups were injected intraperitoneally with 3ng of Diptheria toxin daily for two or three consecutive days for cell ablation experiments. Brains were harvested at P6 or P21 for frozen tissue sections.

### Tissue processing for histological analysis

Brains were retrieved from P0 to P8 after cervical dislocation and from deeply anesthetized adults after transcardial perfusion with 4% paraformaldehyde (PFA). Animals were anesthetized with 100mg/kg ketamine and 5 mg/kg of Xylazine. Brains were then fixed with 4% PFA for 48 hours at 4°C and processed for paraffin embedding. Sections of 5 mm were then collected on glass slides (Fisher) and paraffin was removed using xylene in a 3-wash series (1 × 10 minutes, 2 × 5 minutes). Sections were rehydrated with a descending EtOH series (100% (2x), 95%, 75%, 50%) 3 minutes each followed by two 1-minute washes in H_2_O. Sections were stained with hematoxylin and eosin solution. Golgi stains on adult sections were performed using FD Rapid GolgiStain Kit (FD NeuroTechnologies, Inc) according to manufactural instruction.

### Immunohistochemistry

Immunohistochemistry analyses were performed on frozen tissue sections. Fixed brain tissues described above were rinsed with PBS and immersed in 30% sucrose solution before embedding in tissue freezing medium (OCT). Frozen tissues were sectioned on the Leica CM1950 cryostat at 15 mm. Endogenous peroxidases were blocked with 3% H_2_O_2_ in MeOH (5 mL 30% H_2_O_2_ in 45 mL 100% MeOH for 10 minutes followed with a 1x PBS wash for 3 × 5 minutes. Sections were incubated with PBS blocking solution containing 10% normal goat or donkey serum and 0.1% Triton X-100 at room temperature for 1 hour. A moisture chamber was prepared and slides placed into it. 100 μL of primary antibody (prepared in PBS blocking solution) per slide was added and each was covered with a coverslip and allowed to sit overnight at 4°C. The following day, coverslips were removed and slides were washed in 1x PBS, 3 × 5 minutes each. 100 μL of secondary antibody per slide was added and each was covered with a coverslip and allowed to sit for 1 hour at room temperature. Slides were washed in 1x PBS, 3 × 5 minutes each and were mounted using Fluorosave (Millipore) before imaging. For EdU detection, sections were incubated in PBS with 0.1M CuSO4, 0.1M THPTA, 2mM Azide 488, 1M L-Ascorbic at room temperature for 30min. Sections were washed 3×5minutes in PBS and then incubated with 5 μg/ml DAPI at room temperature for 10 minutes. All fluorescent images were acquired on Leica DMi8 microscope or Zeiss LSM 700 confocal microscope, processed using Leica LAS X or Zeiss Zen software and analyzed using NIH ImageJ.

### Rotarod test

Motor coordination and learning were tested using a commercially available (Harvard Apparaus) accelerating rotarod. The cylinder was 3 cm in diameter and was covered with textured rubber. Mice were confined to a section of the cylinder 5.7 cm wide by grey plastic dividers. The height to fall was 16 cm. Mice were placed on the accelerating rotarod whose speed gradually increased from 4 to 40 rpm over the course of a 300s trial. The time taken for the mouse to fall from the rotating rod was recorded. Mice that fell in less than 15 s were given a second trial. Occasionally, mice clung to the rod, and the whole animal rotated along with it. This behavior was classified as a “rotation,” and the time at which this occurred for the first time on each trial was also recorded for each mouse. Thus the rotarod score was defined as latency to fall or to the first rotation, whichever occurred first. Three sessions were conducted on consecutive days, with three trials per session.

### Gait analysis

Mice were placed on a translucent treadmill at a standstill. The treadmill was then turned on at a speed of 20 cm/s, and still images of their paw movements and placements were taken with a high-speed video camera underneath the belt. Only 10-20 seconds of footage was recorded to obtain ∼40 indices of gait from the analysis software. The following parameters were measured: stance time, time elapsed while foot is in contact with the treadmill, in its stance phase; swing time, time elapsed while foot is in the air, in its swing phase; stride time, time elapsed between two successive initiations of stances; brake time, time elapsed from the first contact with the treadmill to peak of stance; deceleration; propel time, time elapsed from peak stance to full swing.

### Eyeblink conditioning in head-fixed mice

The apparatus and experimental procedure for eyeblink conditioning were previously described (Heiney *et al*, 2014). Briefly, mice were anesthetized with isoflurane (1.5–2% by volume) and positioned on the stereotaxic apparatus prepared for surgery. The skull was exposed, and two small holes were drilled on either side of the midline near bregma for inserting screws. A thin aluminum head plate was then placed over the bregma and the screws were fitted into the central hole in the head plate, which was affixed to the skull by Metabond cement.

Mice were habituated to head restraint for an hour in 3 habituation sesseions on top of a foam cylinder with the head fixed before start of the conditioning sessions. Mice were exposed to either a 240 ms blue LED light positioned 20 cm in front of the mouse or a 240 ms tone of white noise delivered via a speaker (4-Ω magnetic speaker, FF1, TDT) as a conditioned stimulus (CS). The volume of the white noise was set to just below the threshold that causes transient startle movement of the eyelid. The unconditioned stimulus (US) was a periocular air puff (30 – 40 psi) of 40 ms duration and delivered via a 23 gauge needle placed 5 mm from the mouse’s cornea. The pressure of the periocular air puff was set for each mouse to induce a full reflexive blink as the unconditional response (UR). The CS-US inter-stimulus interval was 200 ms. Mice received 100 trials (80 CS-US paired trials and 20 CS-only trials) per day for 14 days. Eyelid movement was detected under infrared illumination using a high-speed (200 or 350 frames/s) monochrome video camera (Allied Vision). Eyelid positions, ranging from 1 (fully closed) to 0 (fully opened), were determined as the fraction of eyelid closure (FEC) using custom MATLAB software and the Video Acquisition Toolbox. The FEC represents the proportion of the distance between the two eyelids. The response was considered to be a conditioned response (CR) if the FEC exceeded 0.2 after the CS onset but before the US.

### Electrophysiology recording

Cerebella from P21 to P30 were retrieved and embedded in agarose while affixed to the specimen holder using adhesive glue. 300 mm thick sagittal slices of cerebellar vermis were prepared using a vibrating microtome (VT1000S, Leica) in oxygenated prechilled medium containing 125 mM NaCl, 2.5 mM KCl, 4 mM MgCl2, 1.25 mM KH2PO4, 26 mM NaHCO3, 1 m CaCl2, and 25 mM D-glucose (pH 7.3–7.4). The slices were then transferred to a recording chamber and superfused with artificial cerebrospinal fluid (aCSF) and gassed with a mixture of 95% O_2_/5% CO_2_ >1 hour at ice-cold temperature before recordings. Whole-cell recordings from Purkinje cells in cerebellar lobules 4-7 (voltage-clamped at −60 mV) were performed at room temperature with borosilicate glass pipettes (2–4 MΩ) pulled with a vertical micropipette puller (PC-10, Narishige). Recording electrodes were filled with internal solution containing 140 mM CsCl, 4 mM NaCl, 0.5 mM CaCl2, 10 mM HEPES, 5 mM EGTA, 2 mM Mg-ATP and 0.4 mM GTP (pH 7.3). For sIPSCs recording, 10 mM 6-Cyano-7-nitroquinoxaline-2,3-dione (CNQX) and 50 to 100 mM amino-5 phosphonopentanoic acid (APV) were added to the external solution to block glutamate receptor-mediated sEPSCs. Membrane currents were recorded using a Multiclamp 700B amplifier (Axon Instruments) connected to a DigiData 1440 (Molecular Devices) using pClamp 10.2 software (Molecular Devices). Series resistance (8–14 MΩ) was monitored throughout the experiments, and experimental data were discarded if the value changed by >20%. All signals were filtered at 2 kHz and sampled at 5–10 kHz. sIPSCs were analyzed with a threshold of 10 pA. Postsynaptic currents were analyzed using Clampfit 11 software (Molecular Devices). Amplitudes and inter-event-interval were measured as a mean of the values obtained from 5 min recording sessions and analyzed using Prism software 8 (GraphPad) for statistical analysis and graphic presentations.

### ERG and VEP Recordings

Mice were dark-adapted overnight, dilated with 1% tropicamide for 10 min, and anesthetized with 20/8/0.8 mg/kg ketamine/xylazine/urethane. Anesthetized mice were placed on the warmed surface of the Celeris ERG system (Diagnosys LLC, Lowell, MA) and corneal electrodes were placed on eyes lubricated with Genteel eye drops. Subdermal platinum needle electrodes were placed in the snout and tail as reference and ground electrodes, respectively. Additional platinum electrodes were placed on the right and left sides of the back of the head, near the visual cortex. The ERG and VEP were recorded sequentially during the same session. For the ERG recording, mice were exposed to 15 flashes of 1Hz, 1cd.s/m^2^ white light. For the VEP recording, mice were exposed to 50 flashes of 1Hz, 0.5 cd.s/m^2^ white light. At the end of the session, mice recovered on a warm pad and then were returned to their cage.

### Quantifications and Statistical Analysis

NIH Image J software was used to measure the area (μm^2^) for regions of interest (ROI) and for the acquisition of cell counting. For each stage, four to six midsagittal sections (approximately 10 um thick) of each cerebellum were used for quantitative analysis. For quantification of cells in the white matter, inner granule layer and molecular layer, the entire laminate was used as ROIs. Cells were counted using Image J plugin Cell counter or Nucleus counter according to the subcellular distribution of target antigen. For quantification of synapses, four to five 5000 μm^2^ rectangular ROIs in the molecular layer of lobule IVand V per sample were used. Positively stained puncta were counted by Image J plugin PunctaAnalyzer. For quantification of PC spine density, DIC images were taken on a Leica TCS SP5 microscope of sagittal cerebellar sections from *Ptf1a*^*Cre*^; *Smo*^*F/-*^ mice and control littermates. Images were taken at a picture size of 1024×1024 pixels, and a 63x objective was used for dendritic spine counting analysis. Z-stacks of individual cells were captured and the length of dendritic spines was measured by tracing a segmented line from the base of the spine to its apex. The number of spines per visible segment was quantified manually in Image J and provided an index of spine density (spines per μm). Manual-counting was done blindly and only spines that had strong, clear Golgi stain were counted. All quantitative data were analyzed using Prism software 8 (GraphPad) for statistical analysis and graphic presentations. Unpaired student’s t-test was used to compare the statistical difference between control and conditional mutant animals.

## Notes

### Competing Interest Statement

The authors have declared no competing interest.

### Summary of Updates

Title revised. Figure 1 revised. Text revised. Supplemental files updated

