## supplementary figures for "Dendritic inhibition by Shh signaling-dependent stellate cell pool is critical for motor learning"

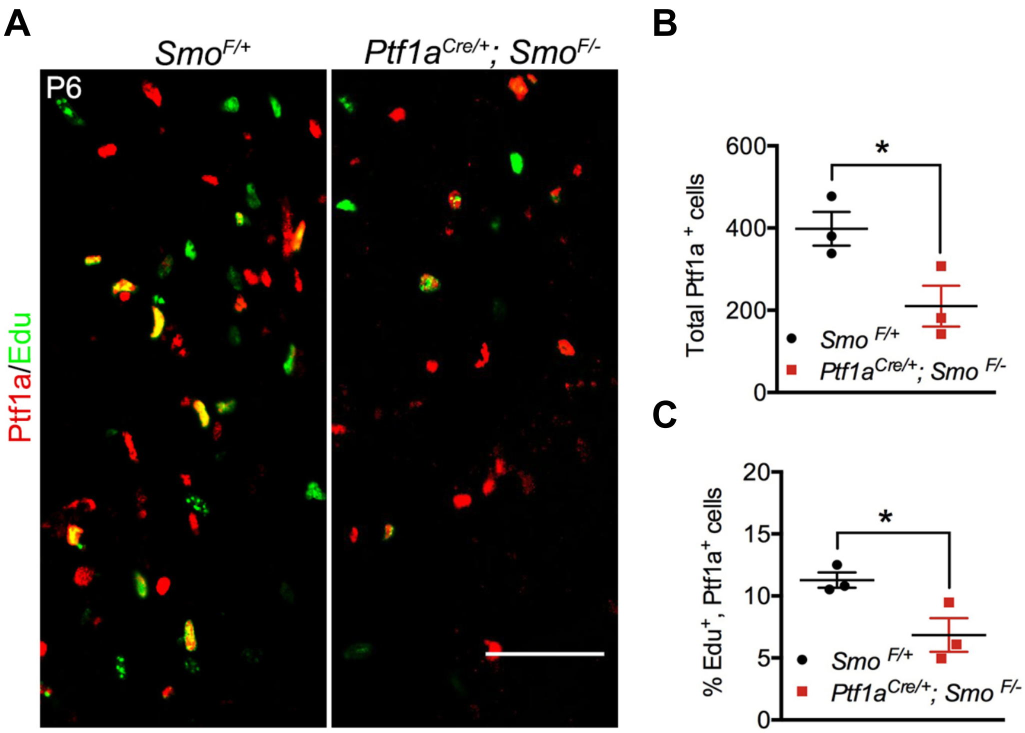


**Figure S1 Shh signaling promotes Ptf1a+ progenitor proliferation** (A-C) Cerebellar parasagittal sections from P6 *Smo* $^{F/+}$ and *Ptf1a*$^{Cre/+}$*; Smo*$^{F/-}$ mice stained with EdU and the antibody against Ptf1a. Quantification of total Ptf1a $^+$ cells in the PWM at P6 (B). The percentage of EdU $^+$/Ptf1a$^+$ double-positive cells relative to the total number of Ptf1a$^+$ cells in PWM at P6 (C) is also quantified. The scale bar represents 50 µm.

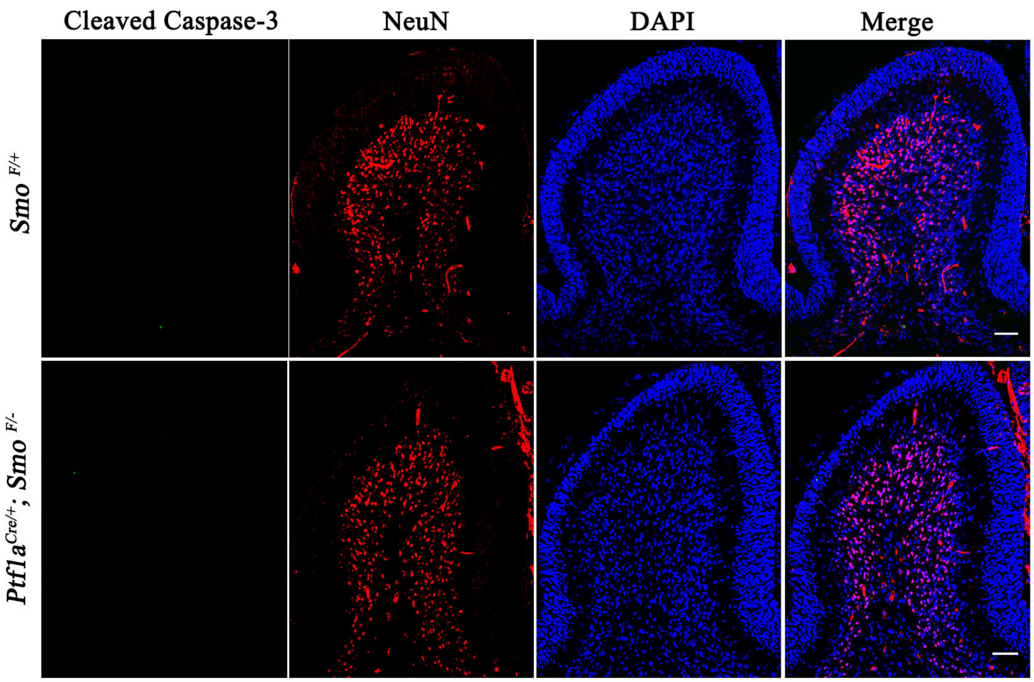


**Figure S2 Loss of Shh signaling in Ptf1a+ progenitors does not activate programmed cell death.**

Cerebellar sections from P6 *Smo* $^{F/+}$ and *Ptf1a*$^{Cre/+}$*; Smo*$^{F/-}$ mice stained with antibodies against cleaved-caspase-3, NeuN and DAPI. The scale bar represents 50 µm.

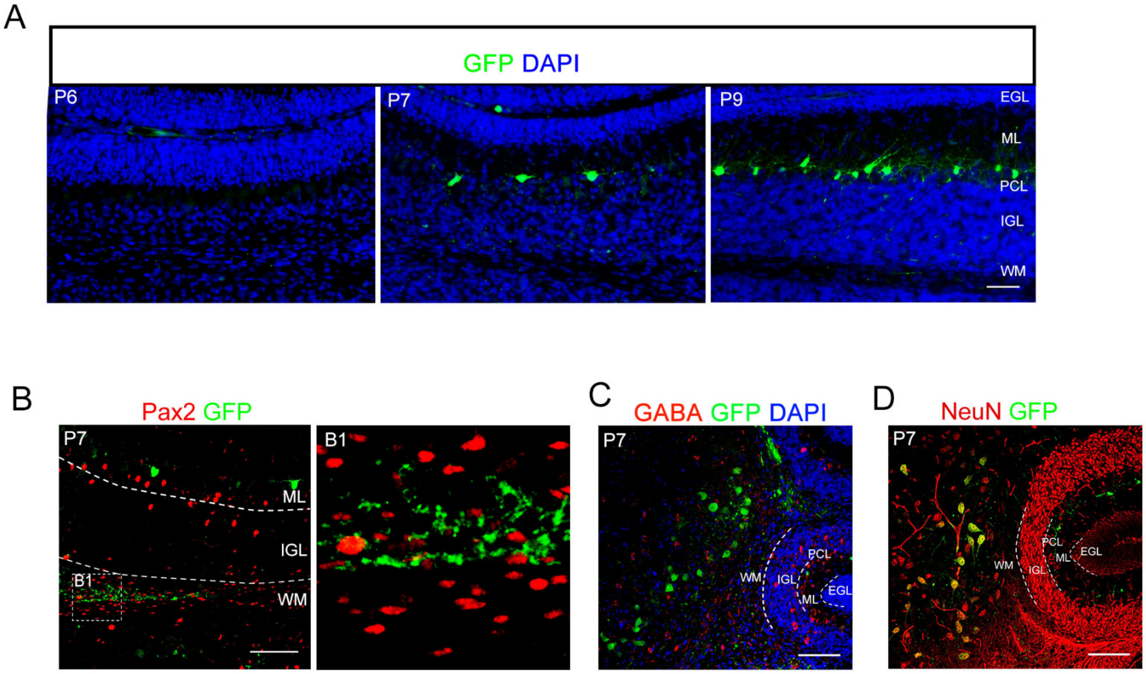


**Figure S3 Ret$^{GFP}$ is selectively expressed in BCs and DCN.**

(A) Expression of Ret$^{GFP}$ in the molecular layer is first detectable at P7.

(B) Ret$^{GFP}$ expression in the PWM is only associated with non-cellular fibers and does not colocalize with Pax2.

(B-C) Ret$^{GFP}$ is also expressed in the NeuN+ DCN (C) and negative for GABA (D).

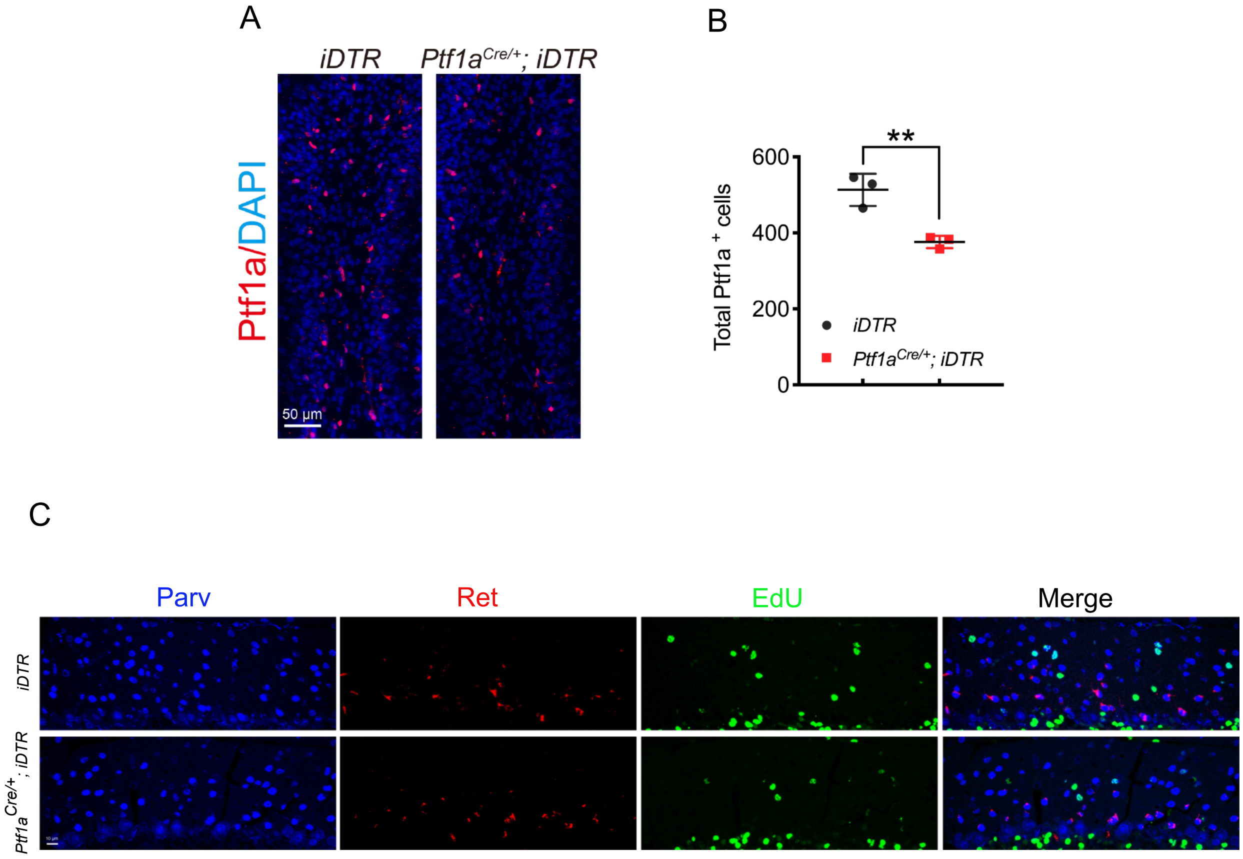


**Figure S4 DT administration efficiently depleted Ptf1a+ progenitors with no delay in BC to SC production.**

(A) Cerebellar sections from P2 *iDTR and Ptf1a*$^{Cre/+}$*; iDTR* mice stained with antibodies against Ptf1a. Scale bars indicate 50 μm.

(B) Quantitative analysis of Ptf1a$^{+}$ cells shown in A. N= 3 mice per group.

(C) Cerebellar sections from P21 *iDTR and Ptf1a*$^{Cre/+}$*; iDTR* mice stained with EdU and antibodies against Parv and Ret.

All graphs displayed are mean ± SEM. * $p < 0.05$. n.s., not significant.

Figure S4, Li et al.

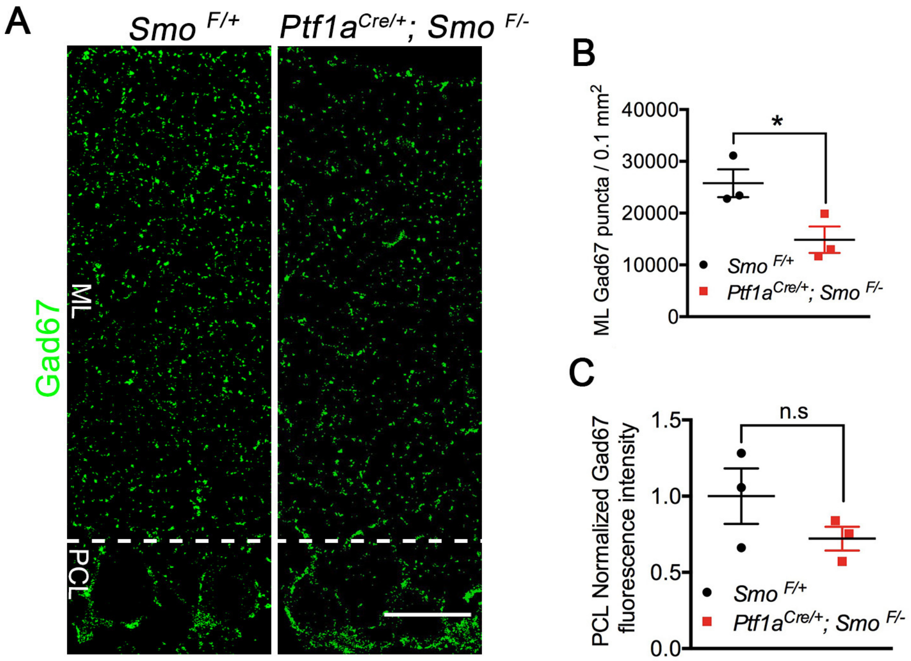


**Figure S5 Loss of Shh-dependent stellate cells reduces ML GABAergic presynaptic puncta.**

(A) Cerebellar sections from P21 *Smo* $^{F/+}$ *and Ptf1a*$^{Cre/+}$*; Smo*$^{F/-}$ mice stained with antibodies against GAD67. Scale bars indicate 25 µm.

(B-C) Quantitative analysis of GAD67 in the ML (B) and PCL (C). N= 3 mice per group.

All graphs displayed are mean ± SEM. * $p < 0.05$. n.s., not significant.

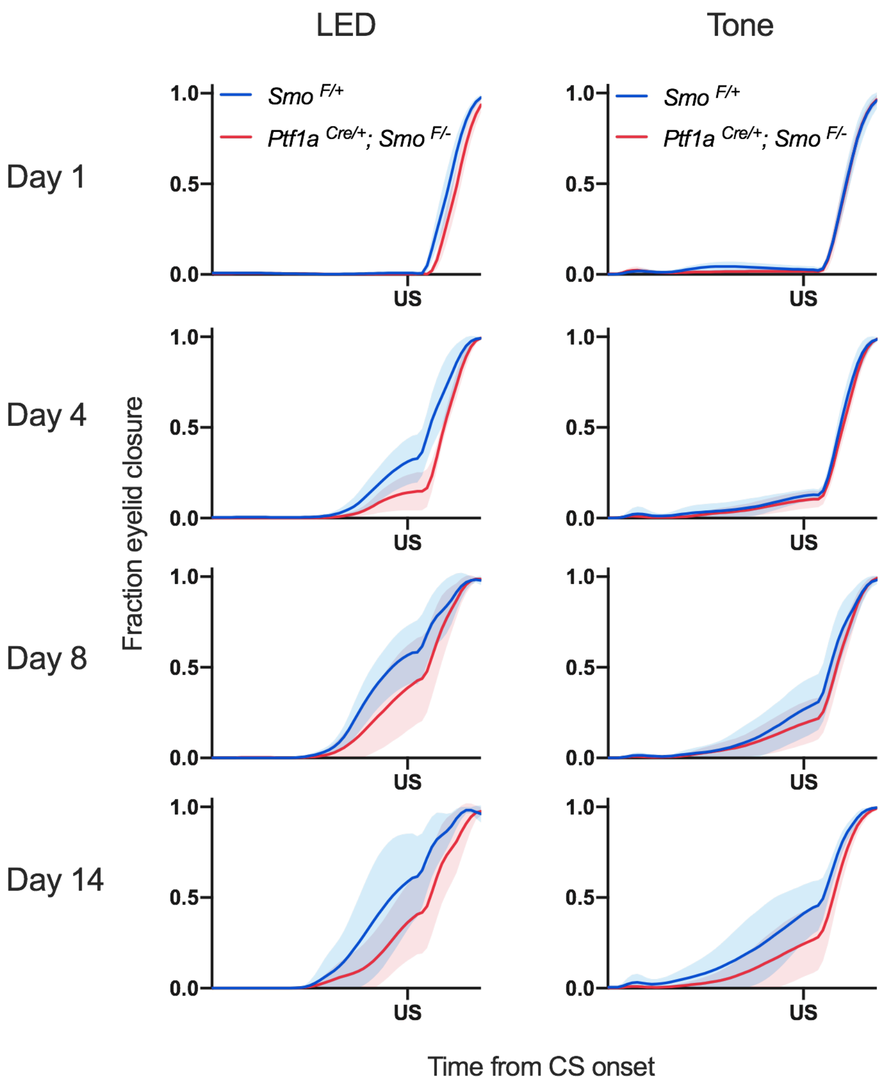


**Figure S6 *Ptf1a*$^{Cre/+}$*; Smo*$^{F/-}$ mice showed the lower amplitude of eyelid closure.**

Average eyelid position traces on Day 1, 4, 8, and 14 for each genotype. Error clouds are standard deviation based on per mouse averages.

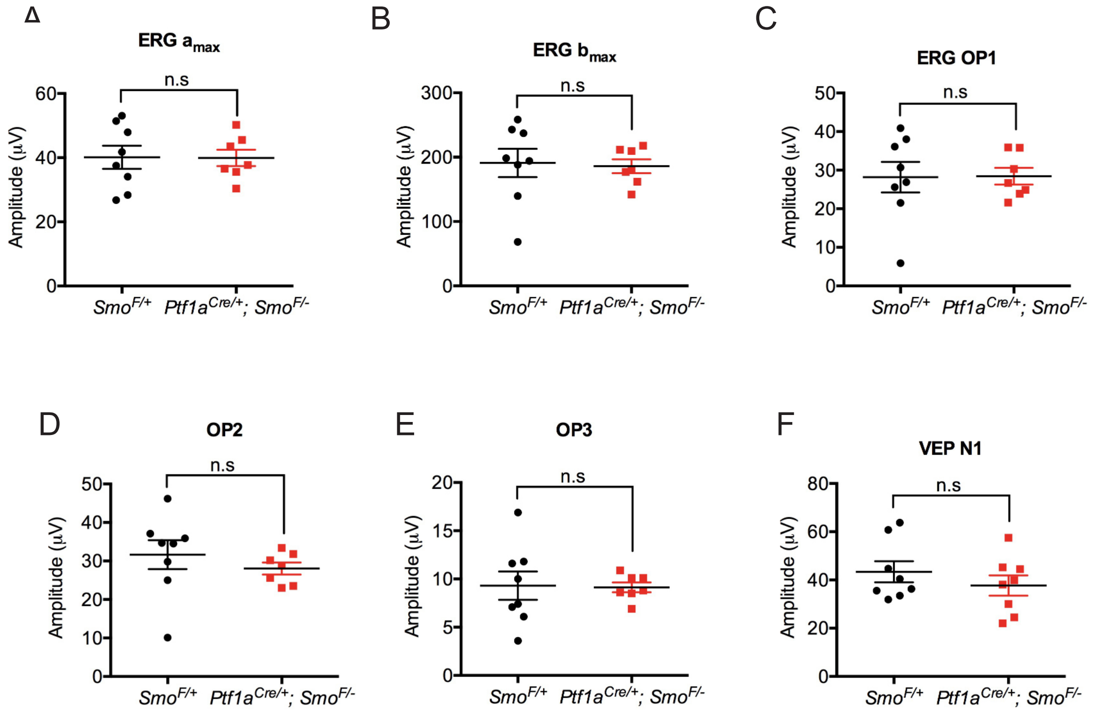


**Figure S7 Normal retinal function in the absence of Shh signaling in Ptf1a+ progenitors.**

(A-F) Statistical analysis of electroretinogram of *Smo* $^{F/+}$ and *Ptf1a*$^{Cre/+}$*; Smo*$^{F/-}$ mice showing $a_{max}$ (A), $b_{max}$ (B), OP1 (C), OP2 (D), OP3 (E), and VEP N1 (F). Values are mean ± SEM. N=8 mice for *Smo* $^{F/+}$ and N= 7 mice for *Ptf1a*$^{Cre/+}$*; Smo*$^{F/-}$. n.s., not significant.

Figure S7, Li et al.

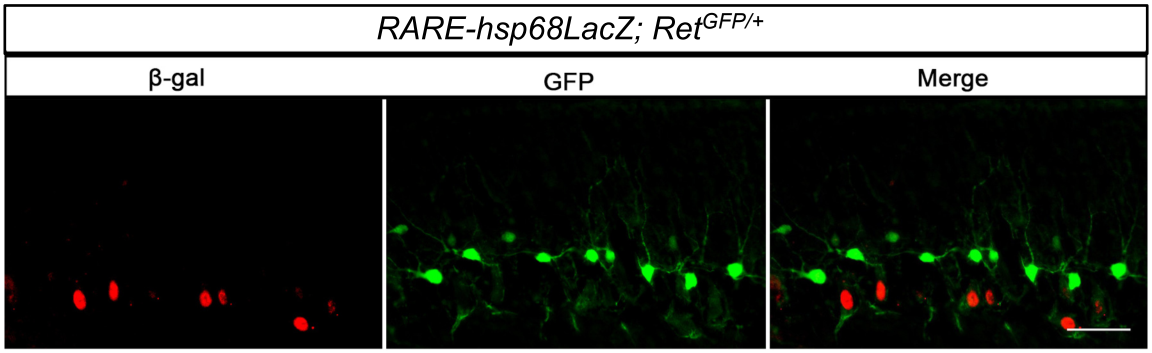


**Figure S8 Retinoic acid signaling is not activated in BCs.**
Cerebellar sections from P21 *RARE-hsp68LacZ*; *Ret*$^{GFP/+}$ mice stained with antibodies against b-gal and GFP.

Figure S8, Li et al.
